# Identification of the m^6^Am methyltransferase PCIF1 reveals the location and functions of m^6^Am in the transcriptome

**DOI:** 10.1101/485862

**Authors:** Konstantinos Boulias, Diana Toczyłowska-Socha, Ben R. Hawley, Noa Liberman-Isakov, Ken Takashima, Sara Zaccara, Théo Guez, Jean-Jacques Vasseur, Françoise Debart, L. Aravind, Samie R. Jaffrey, Eric Lieberman Greer

## Abstract

mRNAs are regulated by nucleotide modifications that influence their cellular fate. Two of the most abundant modified nucleotides are *N*^6^-methyladenosine (m^6^A), found within mRNAs, and *N*^6^,2’-*O*-dimethyladenosine (m^6^Am), which is found at the first-transcribed nucleotide. A long-standing challenge has been distinguishing these similar modifications in transcriptome-wide mapping studies. Here we identify and biochemically characterize, PCIF1, the methyltransferase that generates m^6^Am. We find that PCIF1 binds and is dependent on the m^7^G cap. By depleting PCIF1, we definitively identified m^6^Am sites and generated transcriptome-wide maps that are selective for m^6^Am and m^6^A. We find that m^6^A and m^6^Am misannotations largely arise from mRNA isoforms with alternate transcription-start sites. These isoforms contain m^6^Am that appear to map to “internal” sites, increasing the likelihood of misannotation. Using the new m^6^Am annotations, we find that depleting m^6^Am does not affect mRNA translation but reduces the stability of a subset of m^6^Am-annotated mRNAs. The discovery of PCIF1 and our accurate mapping technique will facilitate future studies to characterize m^6^Am’s function.

## Introduction

An emerging concept in gene expression regulation is that mRNA can be subjected to dynamic and reversible methyl modifications that influence the fate of the transcript in the cell. The vast majority of these regulated methyl modifications occur on two similar nucleotides: adenosine (A) and 2′-*O*-methyladenosine (Am) (Perry et al., 1975; Wei et al., 1975). In the case of adenosine, METTL3 catalyzes the methylation on the N6 position of the adenine ring to form *N*^6^-methyladenosine (m^6^A) (Bokar et al., 1997). Approximately 1-4 m^6^A form in an mRNA on average (Perry et al., 1975), and at least 25% of mRNAs contain at least one m^6^A (Dominissini et al., 2012; Meyer et al., 2012).

N6 methylation also occurs on Am. Am is primarily located at the first transcribed nucleotide position in mRNAs, adjacent to the m^7^G cap. Nucleotides located at the first transcribed nucleotide position in an mRNA are typically methylated on the ribose at the 2′-hydroxyl position. However, if this nucleotide is Am, it can undergo further N6 methylation to form a dimethylated adenosine: *N*^6^, 2′-O-dimethyladenosine (m^6^Am) (Keith et al., 1978; Wei et al., 1975). The m^6^Am-forming methyltransferase is distinct from METTL3 (Keith et al., 1978), but the enzyme that mediates the formation of m^6^Am is currently unknown. However, since m^6^Am is present at the first transcribed nucleotide in ∼30% of all cellular mRNAs, m^6^Am can affect the fate of a large subset of the transcriptome (Wei et al., 1975). Identification of the methyltransferase that regulates m^6^Am will be instrumental for understanding the biological roles of this modification.

Determining whether an mRNA contains m^6^A or m^6^Am, or potentially both modified nucleotides, is important for predicting how epitranscriptomic modification influences the fate of that mRNA in cells. m^6^A is linked to mRNA instability, altered mRNA splicing, translation, and potentially other aspects of mRNA processing (Ke et al., 2015; Ke et al., 2017; Meyer et al., 2015; Sommer et al., 1978; Wang et al., 2015; Xiao et al., 2016). In contrast, m^6^Am has been linked to transcripts that show enhanced mRNA stability and translation (Mauer et al., 2017). Both m^6^A and m^6^Am appear to be dynamic, altered in certain diseases, and susceptible to demethylation (Mauer et al., 2017; Vu et al., 2017; Zhao et al., 2017). Due to the potential for dynamic regulation of m^6^A and m^6^Am levels, it is important to determine if an mRNA of interest contains a modification and to determine whether that modification is m^6^A or m^6^Am.

Currently, transcriptome-wide mapping of m^6^A and m^6^Am uses antibodies that bind 6-methyladenine (6mA). 6mA is the methylated nucleobase that is found in both the m^6^A and m^6^Am nucleotides. The two mapping methods, i.e., MeRIP-Seq (methyl RNA immunoprecipitation followed by sequencing) (Dominissini et al., 2012; Meyer et al., 2012) and miCLIP (m^6^A individual-nucleotide-resolution crosslinking and immunoprecipitation) (Linder et al., 2015) both map sites of 6mA, the methylated nucleobase, rather than m^6^A or m^6^Am. The 6mA “peaks” that are generated by these methods are then interpreted to be either m^6^A or m^6^Am using a variety of criteria. For example, if the 6mA peak is in the 5′ UTR, this suggests that the 6mA peak is caused by m^6^Am since this nucleotide is exclusively found as the transcription-start nucleotides. Nevertheless, it can be difficult to distinguish m^6^Am from m^6^A located within the 5′ UTR of mRNAs. As a result, previous maps of m^6^Am may have inaccuracies which may make it difficult for predicting the function of this modification in mRNA.

To definitively distinguish m^6^A and m^6^Am in transcriptome-wide maps, depletion of either m^6^A or m^6^Am would be required. m^6^A depletion cannot be readily achieved as *Mettl3* is essential for survival in nearly all of 341 cell lines that were screened (Tsherniak et al., 2017). The methyltransferase that generates m^6^Am is not known, but its depletion could enable the identification of the sites that are m^6^Am, since the remaining sites would be m^6^A.

Here we describe the identification of PCIF1 as the methyltransferase that is responsible for generating essentially all m^6^Am residues in mRNA. We show that PCIF1 methylates Am in the context of the m^7^G cap, and has negligible ability to methylate adenosine in RNA outside this context in cells. By mapping 6mA in the transcriptome of PCIF1-deleted cells, we distinguish between m^6^Am and 5’ UTR m^6^A. We find numerous examples where previously annotated m^6^Am sites reflect m^6^A and vice versa. Interestingly, we additionally identified m^6^Am occurring in internal sites relative to reference annotation start sites of numerous genes which were previously identified as m^6^A. The unambiguous mapping of internal m^6^Am nucleosides allows for a precise identification of transcript isoforms with alternative internal transcription-start sites. Characterization of m^6^Am mRNAs in *PCIF1* knockout cells shows that m^6^Am had negligible effects on translation under basal conditions but promotes the stability of a subset of m^6^Am-initiated transcripts. Overall, our studies identify PCIF1 as the methyltransferase that generates m^6^Am in the transcriptome and provides revised transcriptome-wide maps that discriminate between m^6^A and m^6^Am.

## RESULTS

### Identification of PCIF1 as a candidate m^6^Am-forming methyltransferase

Studies in the 1970’s provided initial characterization of an enzymatic activity in HeLa cells that synthesizes m^6^Am (Keith et al., 1978). This activity selectively methylates Am that is adjacent to an m^7^G cap in synthetic RNA substrates (Keith et al., 1978). This activity was shown to be distinct from the methyltransferase that synthesizes m^6^A, now known to be Mettl3 (Bokar et al., 1997). Although the m^6^Am-forming methyltransferase was partially purified, it was not isolated, sequenced or cloned.

In order to identify the m^6^Am-forming enzyme, we performed a comparative bioinformatic analysis of orphan adenosine methyltransferases. These enzymes contain the [DNSH]PP[YFW] motif which is present in all adenine N6-methyltransferases (Iyer et al., 2016). Among these putative adenine methyltransferases, PCIF1 is notable since it evolved at the same time that the 5′ cap emerged in mRNA (Iyer et al., 2016). It has been hypothesized that the 5′ cap has emerged with eukaryotic evolution to replace the Shine-Dalgarno sequence for directing ribosomes to mRNAs and to protect mRNAs from digestion by 5′ exoribonucleases, thus providing an early method for distinguishing self-versus-foreign mRNAs (Furuichi et al., 1977; Shimotohno et al., 1977; Shuman, 2002). Eukaryotes appear to have acquired the progenitor of PCIF1 from an ancestral methyltransferase from the prokaryotic restriction-modification system prior to their divergence from their last common ancestor. The PCIF1 methyltransferase family is derived from the prokaryotic M.EcoKI/M.TaqI methyltransferases of the bacterial restriction-modification systems (Iyer et al., 2016). All of these methyltransferases contain helices before and after the conserved core strand-3 which display partial or complete degeneration into coil elements. Another common feature of these methyltransferases is the addition of a conserved residue from a helix N-terminal to the core methylase catalytic domain. PCIF1 also contains a WW domain which interacts with the C-terminal domain of RNA polymerase II (Fan et al., 2003) (Figure 1A), suggesting that it has a methylation function linked to transcription of mRNA. Based on this, we asked whether PCIF1 is an adenine N6-methyltransferase in mRNA.

**Figure 1.**
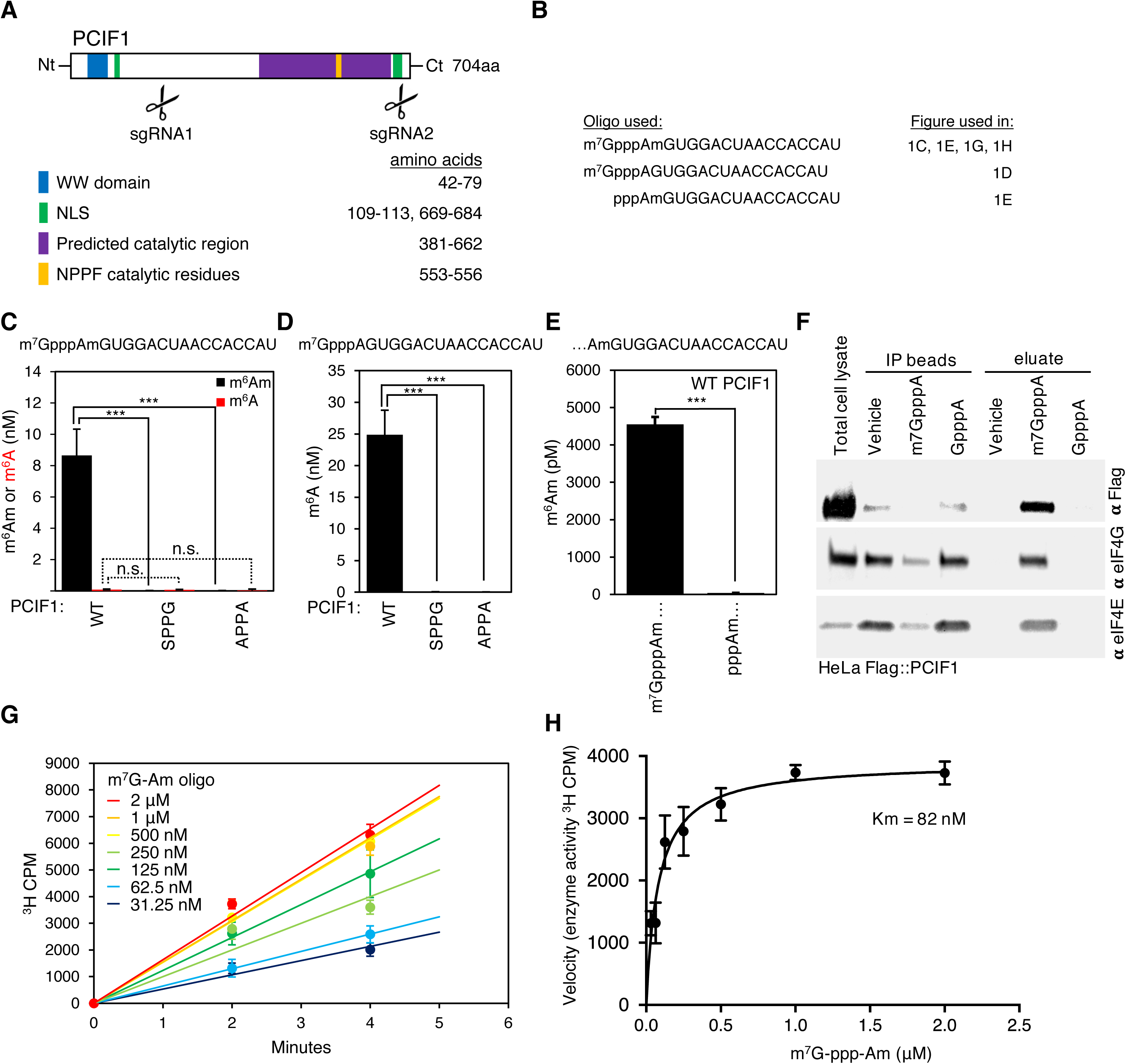
PCIF1 N6-methylates 2′-*O*-methyladenosine *in vitro* in an m^7^G cap-dependent manner. **A.** Schematic of PCIF1 indicating the position of predicted functional domains. The location of the sites of mutations used in the study are shown. The location of the site guide RNAs used in Figure 2 are indicated. **B.** Oligonucleotide sequences used in methyltransferase assays. **C.** PCIF1 methylates synthetic m7G-ppp-Am-N_20_ capped oligonucleotides. GST-PCIF1, but not the catalytically inactive mutants APPA or SPPG efficiently converts m^7^G-ppp-Am to m^7^G-ppp-m^6^Am. A synthetic oligonucleotide m^7^G-ppp-Am-N_20_ (4 μM) was incubated with GST-PCIF1 or the APPA or SPPG PCIF1 mutants (50 nM, 10 min) in the presence of SAM (160 μM), converted Am to m^6^Am, as assessed by UHPLC-MS/MS. Under the same conditions, PCIF1 does not convert any of the 5 internal adenosines to m^6^A, as assessed by UHPLC-MS/MS. Each bar represents the mean ± s.e.m of 3 independent experiments. n.s: not significant, ***: p < 0.001, as assessed by unpaired Student’s t-tests. **D.** PCIF1 can methylate cap-adjacent adenosine regardless of 2′-*O*-ribose methylation. GST-PCIF1, but not the APPA or SPPG PCIF1 mutants efficiently converts m^7^G-ppp-A to m^7^G-ppp-m^6^A. A synthetic oligonucleotide m^7^G-ppp-A-N_20_ (4 μM) was incubated with each enzyme (50 nM, 10 min) in the presence of SAM (160 μM), which converted cap-adjacent adenosine to m^6^A, as assessed by UHPLC-MS/MS. Each bar represents the mean ± s.e.m of 3 independent experiments. ***: p < 0.001, as assessed by unpaired t-tests. **E.** PCIF1 methyltransferase activity *in vitro* toward m^7^G-ppp-Am depends on the presence of the m^7^G cap. A synthetic oligonucleotide m^7^G-ppp-Am-N_20_ (4 μM) or ppp-Am-N_20_ (4 μM) was incubated with GST-PCIF1 (50 nM, 10 min) in the presence of SAM (160 μM), that converted Am to m^6^Am specifically in the m^7^G capped oligonucleotide, as assessed by UHPLC-MS/MS. Each bar represents the mean ± s.e.m of 3 independent experiments. ***: p < 0.001, as assessed by unpaired t-tests. **F.** PCIF1 directly binds the m^7^G cap. Western blot analysis with an anti-FLAG antibody was performed to examine the binding of 3xFLAG-PCIF1 from HeLa cell extracts with m^7^GTP-conjugated beads. The beads were eluted with vehicle, an m^7^GpppA analog competitor, or a non competitor GpppA analog. eIF4E and eIF4G were used to control for binding to m^7^G. Binding was evaluated with Flag, eIF4E, and eIF4G-specific antibodies. **G.** Reaction curves for PCIF1 toward various concentrations of m^7^G-ppp-Am substrates. An oligonucleotide m^7^G-ppp-Am-N_20_ (at indicated concentration) was incubated with GST-PCIF1 (20 nM) for the indicated times in the presence of 1.33 μM ^3^H-SAM and 10 μM SAM. Enzyme activity was determined by the presence of ^3^H signal in the purified oligonucleotide, as assessed by scintillation counting. Each point represents the mean ± s.e.m of 3 independent experiments. **H.** Michaelis-Menten kinetics of PCIF1 N6-methylytransferase activity toward m^7^G-ppp-Am as determined from the reaction curves of Fig. 1G. Each point represents the mean ± s.e.m of 3 independent experiments.

### PCIF1 N6-methylates 2′-*O*-methyladenosine in an m^7^G cap-dependent manner *in vitro*

To identify any potential PCIF1-dependent nucleotide methyltransferase activity, we bacterially expressed and purified glutathione S-transferase (GST)-tagged PCIF1 and performed *in vitro* methyltransferase assays using synthetic RNA oligonucleotides.

To test whether PCIF1 can methylate the cap-adjacent adenosine of mRNAs, we performed *in vitro* methyltransferase assays with purified GST-PCIF1 protein and an RNA oligonucleotide with a 5′ m^7^G cap followed by 2′-*O*-methyladenosine (m^7^G-ppp-Am-N_20_) (Figure 1B). We found that wild-type PCIF1 methylates Am to produce m^6^Am, as assessed by UHPLC-ms/ms (Figure 1C).

Interestingly, we did not detect any m^6^A in these methylation reactions despite the presence of 5 additional internal adenosines in the oligonucleotide sequence (Figure 1C), suggesting that PCIF1 preferentially N6-methylates 2′-*O*-methyladenosine rather than internal adenosines.

To determine whether this methyltransferase activity was intrinsic to PCIF1 and not a contaminating protein, we mutated two critical residues in the catalytic domain of PCIF1 (Figure 1A). PCIF1’s predicted catalytic domain consists of a four amino acid motif, NPPF, where N6-adenine methylation should be regulated by the polar group of the asparagine residue, the following two prolines which prime the NH_2_ group of the target nucleotide for methylation, and an aromatic phenylalanine residue which is predicted to hold the target base in place by a π-π stacking interaction (Iyer et al., 2016). We mutated both asparagine 553 and phenylalanine 556 to alanines (NPPF→APPA) or to a serine and a glycine (NPPF→SPPG) as these mutations were shown to inactivate the N6-methyltransferases EcoKI and Dam while still retaining their protein structure (Guyot et al., 1993; Willcock et al., 1994). We found that neither the APPA nor SPPG mutant was able to methylate the RNA oligonucleotides (Figure 1C), suggesting that PCIF1 possesses Am methyltransferase activity *in vitro*.

To further explore the specificity of PCIF1 methyltransferase activity, we performed *in vitro* methylation assays as above using an RNA oligonucleotide substrate with a 5′ m^7^G cap followed by adenosine (m^7^G-ppp-A-N_20_) rather than Am. We found that wild-type PCIF1 but not the SPPG or APPA PCIF1 mutant was able to N6-methylate adenosine to m^6^A (Figure 1D), suggesting that PCIF1 has the ability to methylate the m^7^G-adjacent adenosine regardless of its 2′-O-methylation status.

Previous characterization of the m^6^Am-forming methyltransferase found that only m^7^G-capped mRNAs were optimal substrates (Keith et al., 1978). To determine if PCIF1 exhibits this property, we performed methyltransferase assays using identical oligonucleotides capped with either m^7^G and a triphosphate bridge (m^7^G-ppp-Am-N_20_) or lacking the m^7^G cap (ppp-Am-N_20_). We found that PCIF1 efficiently methylated Am to m^6^Am in the m^7^G capped oligonucleotide but was unable to methylate Am in the oligonucleotide that lacks the m^7^G cap (Figure 1E), suggesting that PCIF1 methyltransferase activity towards Am depends on the presence of the m^7^G cap.

Because PCIF1 methylates oligonucleotides that are m^7^G capped we wanted to determine whether PCIF1 could bind to the m^7^G cap directly. To test this, we performed cap-binding assays with PCIF1 using 7-methylguanosine-5-triphosphate (m^7^G-ppp)-coupled Sepharose beads. In these experiments, we used lysates from HeLa cells expressing FLAG-tagged wild-type PCIF1. As had been previously reported (Sonenberg et al., 1978; Sonenberg and Shatkin, 1977), we found that cap-binding proteins eIF4E and eIF4G were bound to m^7^G-ppp beads and were efficiently eluted using an m^7^GpppA competitor but not a GpppA competitor (Figure 1F). Similarly, we found that PCIF1 bound to the m^7^G-ppp beads and was eluted with an m^7^GpppA competitor but not with a GpppA competitor (Figure 1F). Together, these data suggest that PCIF1 binds directly to the m^7^G cap, which may account for its specificity towards adenosine adjacent to the m^7^G.

To assess the rate at which PCIF1 N6-methylates the m^7^G-adjacent Am we performed methyltransferase assays using a serial dilution of the m^7^GpppAm capped oligonucleotide, PCIF1, and tritiated *S*-adenosyl methionine [^3^H]-SAM as the methyl donor (Figure 1G). A Michaelis-Menten analysis yielded a K_M_ = 82 +/-18.23 nM (Figure 1H). A similar K_M_ was reported for the m^6^A mRNA methyltransferase complex Mettl3/Mettl14 (22-102 nM, (Li et al., 2016)) and the DNA 5mC methyltransferases DNMT enzymes (19-329 nM, (Hemeon et al., 2011; Robertson et al., 2004)) and lower K_M_ was reported for the m^7^G mRNA cap methyltransferase RNMT (210 nM, (Martin and Moss, 1976)) or the 2′-hydroxyl mRNA ribose methyltransferase CMTR1 (1 µM, (Belanger et al., 2010)). Together this data demonstrates that PCIF1 has robust N6-methyltransferase activity toward the 2′-*O*-methyladenosine adjacent to m^7^G in RNA.

### *PCIF1* knockout abolishes m^6^Am levels without affecting m^6^A in RNA

To determine the ability of PCIF1 to generate m^6^Am in cells, we used CRISPR to delete *PCIF1* in various cell lines (Figures 2A and **S1A**). We then examined levels of m^6^Am and m^6^A in RNA in these *PCIF1* knockout cells.

**Figure 2.**
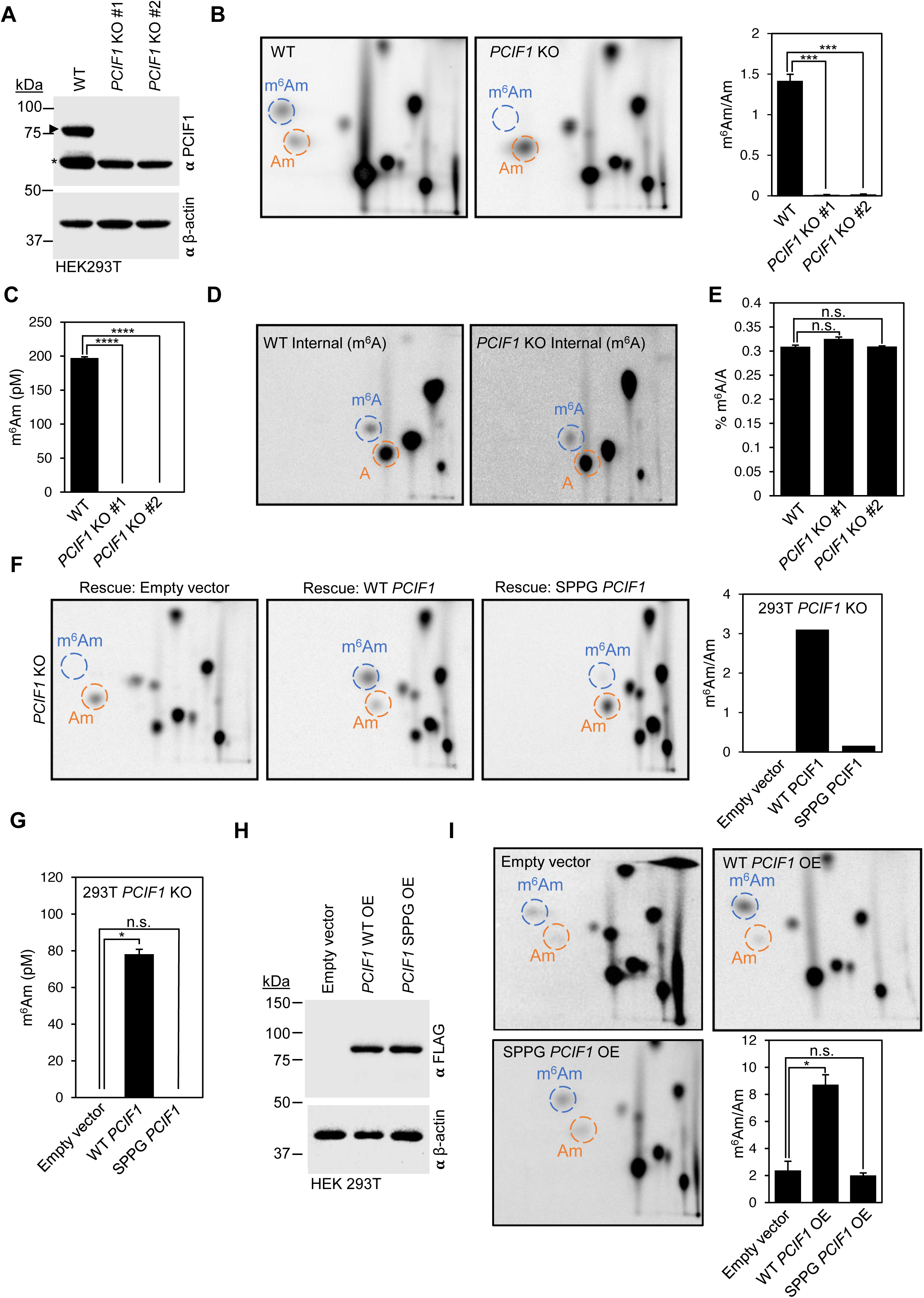
PCIF1 N6-methylates 2′-*O*-methyladenosine in cells. **A.** CRISPR-mediated *PCIF1* knockout in HEK293T cells was assessed by western blot of whole cell extracts using an anti-PCIF1 antibody. The upper band represents endogenous PCIF1, whereas the lower band is a non-specific band. β-actin is shown as a loading control. **B.** PCIF1 is required for formation of m^6^Am in mRNA in cells. Poly(A) RNA was purified from *PCIF1* knockout HEK293T cells and m^6^Am and Am levels were quantified by 2D-TLC. In this method, poly(A) RNA was decapped and the 5′ nucleotide was selectively radiolabeled by 5′ phosphorylation. After RNA hydrolysis, the radiolabeled 5′ nucleotide was detected and quantified based on the position (indicated) of m^6^Am and Am on 2D-TLC. The m^6^Am spot was lost in *PCIF1* knockout cells relative to wild-type (WT) cells. Representative images are shown from 3 biological replicates. The blue circles representing m^6^Am and the orange circles representing Am as assessed by standards run on separate blots in parallel. The bar graph on the right represents the mean ± s.e.m of 3 independent experiments. *** p < 0.001 as assessed by Student’s t-test. **C.** m^6^Am is depleted in *PCIF1* KO HEK293T cells as assessed by UHPLC-MS/MS. Each bar represents the mean ± s.e.m of 3 independent experiments. ****: p <0.0001 as assessed by paired t-tests. **D.** PCIF1 does not affect the level of internal m^6^A. 2D-TLC analysis of poly(A) RNA from HEK293T (WT) and *PCIF1* KO HEK293T show no effect on the level of m^6^A. **E.** Internal m^6^A is not affected by PCIF1 deletion in HEK293T cells as assessed by UHPLC-MS/MS. Each bar represents the mean ± s.e.m of 3 independent experiments. Ns: not significant, as assessed by paired t-tests. **F.** Wild-type but not a catalytically inactive PCIF1 mutant can restore m^6^Am levels in *PCIF1* knockout cells as assessed by 2D-TLC. Relative abundance of modified adenosines in mRNA caps of HEK293T KO control cells (Empty Vector) and *PCIF1* knockout cells stably expressing 3X-FLAG-PCIF1 WT or 3X-FLAG-PCIF1 SPPG. **G.** Wild-type but not catalytically inactive PCIF1 mutant can restore m^6^Am levels in *PCIF1* KO cells as assessed by UHPLC-MS/MS. Each bar represents the mean ± s.e.m of two independent experiments. Ns: not significant, *: p < 0.05 as assessed by paired t-tests. **H.** Western blot analysis demonstrate equivalent expression of wild-type and catalytically inactive PCIF1 in HEK293T cells. Whole cell extracts from HEK293T control cells (Empty Vector) and HEK293T cells stably overexpressing 3X-FLAG-PCIF1 WT or 3X-FLAG-PCIF1 SPPG catalytically inactive mutant were blotted with anti-FLAG antibody. β-actin is used as a loading control. **I.** Overexpression of wild-type but not catalytic mutant PCIF1 increases m^6^Am levels in HEK293T cells as assessed by 2D-TLC. Upper and left panels show representative images of 3 independent experiments with the blue circles representing m^6^Am and the orange circles representing Am as assessed by standards run on separate blots in parallel. The bar graph represents the mean ± s.e.m of 3 independent experiments. Ns: not significant, \*\*\*\**P* ≤ 0.0001, as assessed by unpaired t-tests.

To measure m^6^Am, we used a two-dimensional thin-layer chromatography (2D-TLC)-based method that can measure both m^6^Am and Am, allowing the ratio of these modified forms of adenosine to be readily detected in poly(A) RNA (Kruse et al., 2011). In this assay, mRNA is decapped, and the 5′ nucleotide in RNA is selectively radiolabeled with [^32^P]-ATP by polynucleotide kinase (PNK). In this way, the first transcribed nucleotide in the transcriptome can be selectively detected and quantified. As expected, all the known nucleotides located at the first transcribed nucleotide in poly(A) RNA were detected, i.e., m^6^Am, Am, Gm, Cm, and Um. However, in *PCIF1* knockout cells, a selective and complete loss of m^6^Am was detected (Figure 2B). Thus, PCIF1 is required for the presence of m^6^Am at the first transcribed nucleotide in mRNA.

Similarly, we were unable to detect m^6^Am in mRNA of *PCIF1* knockout cells mRNA as assessed by UHPLC-ms/ms (Figure 2C). This dramatic effect on m^6^Am levels was not restricted to HEK293T cells as we observed a similar effect in HeLa cells lacking PCIF1 (Figure S1B).

To determine if PCIF1 exhibited N6-methyltransferase activity against internal adenosines, we asked if PCIF1 deletion affects m^6^A levels in mRNA. To test this, we first used a 2D-TLC-based method that selectively detects m^6^A in the G-A-C context in mRNA (Zhong et al., 2008). m^6^A was readily detected in poly(A) RNA in control cells, and no reduction was seen in *PCIF1* knockout cells (Figure 2D).

To confirm these results, we measured m^6^A levels in poly(A) RNA using UHPLC-MS/MS. We found no change in m^6^A levels in *PCIF1* knockout cells relative to wild-type cells (Figure 2E).

To confirm that the loss of m^6^Am in the CRISPR knockout cells was due to a loss of PCIF1 itself, we performed rescue experiments. In these experiments, we used wild-type or the SPPG catalytically inactive PCIF1 mutant (Figure S1C). We found that re-expression of the wild-type but not the catalytically inactive PCIF1 could restore m^6^Am levels in mRNA of HEK293T *PCIF1* knockout cells as assessed by 2D-TLC (Figure 2F) and by UHPLC-ms/ms (Figure 2G).

The inability of catalytic inactive PCIF1 mutant to rescue the effects of PCIF1 deletion was not due to a misexpression or mislocalization of the mutant protein, as both wild-type and catalytically inactive PCIF1 were expressed at similar levels (Figure S1C) and were localized predominantly to the nucleus (Figure S1D), which is in agreement with the previously reported nuclear localization of PCIF1 (Hirose et al., 2008). Together, these data indicate that PCIF1 is required to form essentially all m^6^Am in mRNA.

To test whether PCIF1 was sufficient to increase m^6^Am levels in cells, we measured m^6^Am levels in mRNA upon overexpressing PCIF1. We found that PCIF1 overexpression in HEK293T cells (Figure 2H) led to a ∼3-fold increase in the m^6^Am to Am ratio (Figure 2I). This increase in m^6^Am levels was dependent on the catalytic activity of PCIF1, as overexpression of a catalytically inactive PCIF1 mutant had no effect on m^6^Am levels (Figure 2I). Together this data suggests that PCIF1 is both necessary and sufficient to methylate m^6^Am on mRNA in cells.

### miCLIP analysis of *PCIF1* knockout cells distinguishes m^6^Am from 5′ UTR m^6^A residues

Next, we used the *PCIF1* knockout cells to distinguish between m^6^Am and m^6^A in transcriptome-wide 6mA maps. In these experiments, we used the miCLIP method, a protocol that produces narrow peaks, and nucleotide transitions at and adjacent to the m^6^A (Linder et al., 2015). m^6^A is nearly universally followed by cytosine in mRNA (Wei et al., 1976). This C is frequently observed to undergo a C to T transitions as a result of antibody crosslinking in miCLIP, which can then be used to identify m^6^A (Linder et al., 2015). Although C to T transitions are useful for detecting m^6^A, m^6^Am can also be followed by cytosine. Thus, transitions alone are not sufficient to distinguish between m^6^A and m^6^Am. To identify m^6^Am sites, the peak shape has been used (Linder et al., 2015). A peak caused by m^6^Am should exhibit a unique peak shape that exhibits a marked drop off of reads at an annotated A-starting transcription-start site. However, because m^6^A close to the transcription-start site would also produce a similar drop-off of reads, this approach may result in false positive m^6^Am identifications.

Furthermore, these approaches are highly dependent on transcript annotations that may not have accurate transcription-start site information for the cell type investigated. For example, RefSeq and ENSEMBL frequently show different annotations for the transcription-start site for the same gene (Zhao and Zhang, 2015). As such, many true m^6^Am peaks may have been discarded or thought to be m^6^A based on their location away from a transcription-start site.

We therefore used *PCIF1* knockout cells to distinguish m^6^Am and m^6^A. We mapped 6mA peaks using miCLIP in control and *PCIF1* knockout cells. In control cells, the overall distribution of reads shows a marked enrichment of reads in the vicinity of the stop codon as well as the transcription-start site, which is generally assumed to reflect m^6^A and m^6^Am, respectively (Figure 3A). The *PCIF1* knockout cells exhibited a clear drop in reads that map near the annotated transcription start site (Figure 3A), suggesting these reads derive from an m^6^Am residue.

**Figure 3.**
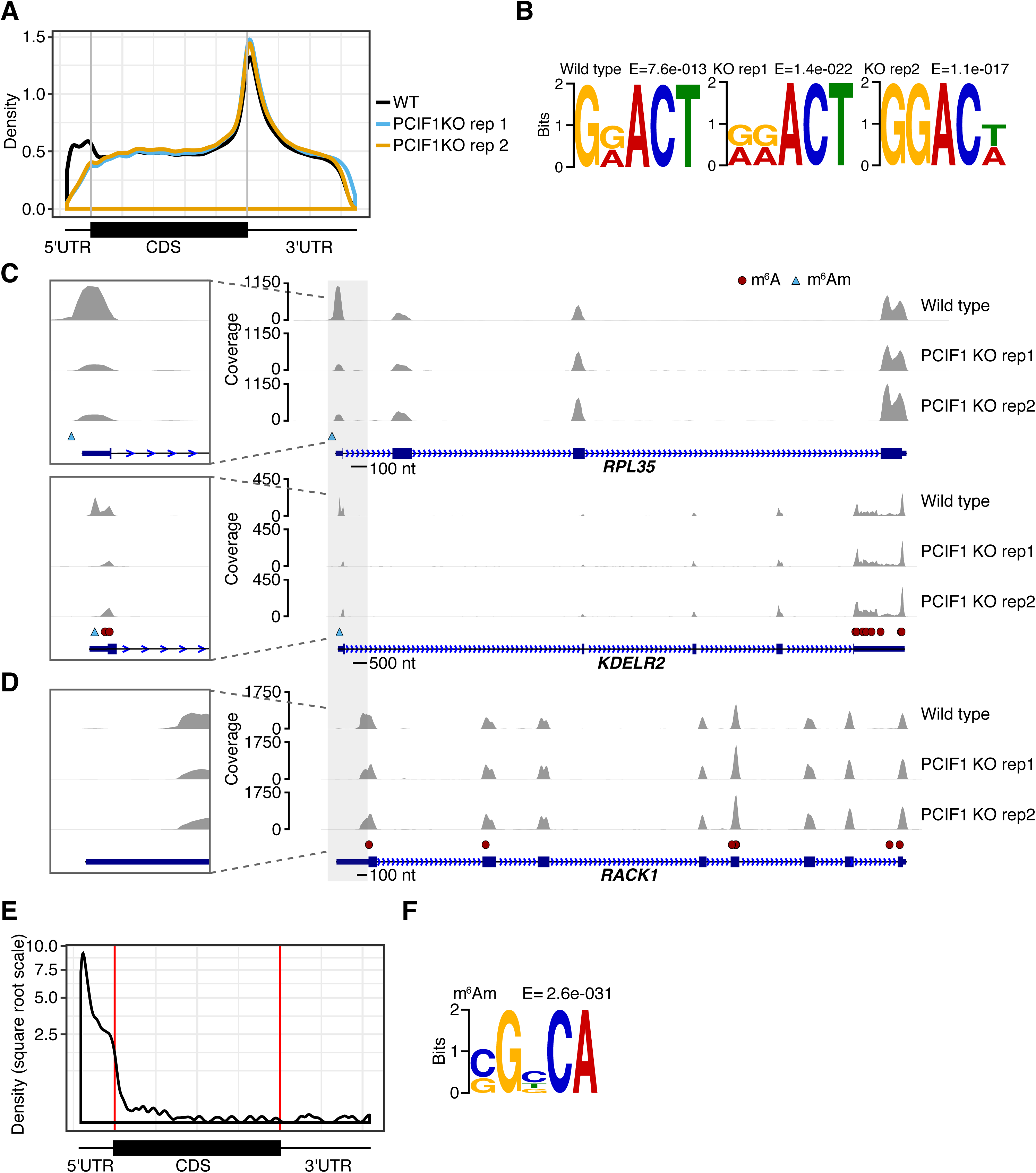
Depletion of PCIF1 distinguishes m^6^A and m^6^Am in transcriptome-wide 6mA maps. **A.** Metagene of miCLIP reads in wild-type and *PCIF1* knockout HEK293T cells. Shown is a metagene analysis of each read obtained from either the wild-type or PCIF1 knockout (KO) miCLIP dataset. The first nucleotide of each read (with respect to the RNA strand) was extracted and plotted. The relative positions of the nucleotide within the transcript body was determined using the longest RefSeq-annotated transcript isoform for each gene using MetaPlotR package (Olarerin-George and Jaffrey, 2017). Comparison between wild type (black) and *PCIF1* knockout (blue and orange) show an overall highly similar profile with read enrichment near the stop codon. However, the enrichment of reads in the 5′ UTR of the wild-type miCLIP was lost in the *PCIF1* knockout dataset, suggesting a complete loss of m^6^Am in the *PCIF1* knockout cells. **B.** Motif search within called peaks show the DRACH motif as the most enriched in all datasets. DREME was used to identify short, ungapped motifs in called peaks in each condition. The top enriched motif by E-value is shown. In each case, the classic m^6^A consensus motif was discovered, consistent with the idea that m^6^A is the most abundant 6mA-containing nucleotide mapped by miCLIP. **C.** miCLIP peaks can be identified as m^6^Am or m^6^A based on their decrease in *PCIF1* knockout cells. Genome tracks were plotted for *RPL35* and *KDELR2* with called m^6^A sites (FDR<0.1) and m^6^Am sites indicated by red circles and blue triangle, respectively. Zoomed insets also show this dataset can distinguish between an m^6^Am peak and close-by m^6^A sites. Scale bars are shown underneath gene model. **D.** The previously annotated m^6^Am site in *RACK1* is actually a 5′ UTR m^6^A. The transcription-start site-proximal peak in *RACK1* is not affected in the *PCIF1* knockout. Inspection of the miCLIP peak shows overlap with a DRACH motif, further supporting the idea that this peak derives from a m^6^A residue. Scale bar is shown underneath gene model. **E.** Metagene analysis of *PCIF1* knockout-validated m^6^Am sites shows m^6^Am sites throughout the 5′ UTR and in the transcript body. Shown is a metagene of the exact sites of m^6^Am within the PCIF1-dependent peaks as determined by A to T transitions and the read drop-off method. The metagene reveals an overall enrichment at the transcription-start site, with some sites that appear to be within the CDS and 3′ UTR. **F.** The m^6^Am transcription-start nucleotide is found in a genomic BCA context. As in Figure 3B, a DREME motif search of the nucleotides surrounding each m^6^Am was performed. This confirms the previously reported BCA motif, and shows that the promoter sequence upstream of the m^6^Am is GC-enriched.

A motif analysis of significant peaks showed the DRACH m^6^A consensus (D = A, G, U; R = A, G; H = A, C, U) as the most common motif in each data set (Figure 3B).

We next examined the 6mA peaks in transcripts that showed differences in the control and *PCIF1* knockout miCLIP datasets. As expected, we were able to detect a loss of peaks near the transcription-start site of certain genes in the *PCIF1* knockout. For example, *RPL35* and *KDELR2* show peaks near the annotated transcription-start site as well as at internal sites (Figure 3C). The transcription-start site-proximal peaks were completely lost in the *PCIF1* knockout miCLIP dataset. These data are consistent with the idea that the transcription-start site peaks reflect m^6^Am.

However, in some cases, the peaks near the transcription-start site were not affected in the *PCIF1-*knockout dataset. For example, peaks are readily detectable near the transcription-start sites of *RACK1* and *RPS5* and were previously annotated as m^6^Am in HEK293T cells based on peak shape and lack of C to T transitions (Mauer et al., 2017). However, these peaks persist in the *PCIF1* knockout dataset (Figure 3D and Figure S2A). These peaks overlap a DRACH consensus, and in this HEK293T miCLIP dataset C to T transitions are detected for *RACK1* (Figure 3D), suggesting that these sites are actually m^6^A.

The variability in C to T transitions reflects the low transition rate induced by the antibody adduct on this transcript. Overall, these data indicate that PCIF1 depletion can be used to determine the identity of an m^6^A peak.

Overall, only 60.2% of genes that had previously annotated as m^6^Am (Mauer et al., 2017) were validated as m^6^Am based on their loss in *PCIF1* knockout cells. In some cases, this could be explained by peaks being below the threshold for detection in one or both replicates. Nevertheless, this difference highlights the importance of depleting the modification writer to prevent false positives.

### A high-confidence transcriptome-wide map of m^6^A and m^6^Am based on PCIF1 depletion

We next wanted to create a high confidence map of all m^6^A and m^6^Am sites in the transcriptome. To map m^6^Am, we searched for all peaks that exhibit a marked reduction in miCLIP signal in the *PCIF1* knockout dataset. The majority of peaks, which likely reflect m^6^A, showed no substantial difference between control and *PCIF1* knockout miCLIP datasets (Figure S2B). However, this search identified 2360 peaks which exhibited a significant reduction in both *PCIF1* knockout datasets (Figure S2B, p = 0 by hypergeometric probability).

We next identified the exact m^6^Am residue within each of these peaks. In our previous approach, we used a “pile up” of reads that drop off at the 5′ end of these read clusters in A-starting genes to predict the m^6^Am site (Linder et al., 2015). In some cases, the drop off is not easily detected or several of these were found in close proximity. This appears to occur when (1) the total reads is too few; or (2) the reads terminate before the transcription-start site, possibly due to impaired reverse transcription through the 2′-*O*-methyl modifications (Maden et al., 1995) in the cap-proximal nucleotides, or due to non-templated nucleotide addition that occurs at the ends of cDNAs generated by reverse transcriptases (Chen and Patton, 2001).

Therefore, we wanted to develop an alternative approach to identify m^6^Am within the PCIF1-dependent peaks. Previously we showed that antibodies can induce A to T transitions at the m^6^A site in miCLIP (Linder et al., 2015). This is readily detected at methylated, but not nonmethylated, adenines throughout the transcriptome (Figure S2C) including transcription-start sites and near stop codons, reflecting m^6^Am and m^6^A, respectively (Figure S2D).

Therefore, we used a 10% A to T transition rate to identify the m^6^Am site within PCIF1-dependent peaks. The drop-off approach was used when the A to T transition rate did not meet these criteria (Figure S2E). However, it should be noted that there was high similarity in the m^6^Am sites that were called when using these methods separately (Figure S2F, p= 3.4e-04 by hypergeometric probability).

Overall, the m^6^Am sites mapped based on their dependence on PCIF1 (**Table S1**) were primarily located throughout the 5′ UTR, with a prominent enrichment at the annotated transcription-start site and a marked reduction in the frequency of sites at the start codon (Figure 3E). Motif analysis of the genomic context of the exact m^6^Am nucleotide revealed the BCA motif as was previously reported (Linder et al., 2015); additionally, this shows the upstream promoter sequence is GC-enriched (Figure 3F).

Next, we mapped m^6^A in the 5′ UTR. As in the miCLIP protocol (Linder et al., 2015), significantly enriched C to T transitions in a DRACH consensus were used to call m^6^A sites. This identified 399 5′ UTR m^6^A sites that were robustly called across all datasets (**Table S2**).

We next asked if 5′ UTR m^6^A and m^6^Am have distinct functions, based on the updated m^6^A and m^6^Am sites called here. Functional annotation analysis using DAVID shows that transcripts containing these distinct modified nucleotides are linked to different functions, with 5′ UTR m^6^A associated with processes such as transcription and cell division, while m^6^Am is primarily associated with splicing (Figure S2G,H and **Table S3**).

### *ATF4* contains a m^6^Am rather than m^6^A in its 5′ UTR

A particularly prominent 5′ UTR m^6^A site has been described in *ATF4*, which has been described as mediating its unusual stress-regulated translation (Zhou et al., 2018). *ATF4* has two upstream open reading frames (uORFs) in the 5′ UTR, and due to their translation the main open reading frame, which encodes the ATF4 protein, is not translated under basal conditions in human and mouse cells (Vattem and Wek, 2004). However, during stress, the second uORF is skipped, and the ribosome scans to the main open read frame after translating the first uORF. This allows the ATF4 protein to only be translated during stress. m^6^A was mapped to the second open reading frame and was described as disappearing in a stress-dependent manner, thus mediating the ability of ATF4 translation to switch from the second uORF to the main open reading from during stress (Zhou et al., 2018).

However, using miCLIP it is apparent that the 6mA peak in the 5′ UTR of *ATF4* is not located within the second open reading frame (Figure S3A). Instead the peak is located at the transcription-start nucleotide and does not overlap with the position of the putative m^6^A.

Based on the location of the peak, we asked if it instead reflects m^6^Am rather than m^6^A. To test this, we examined *ATF4* in the *PCIF1* knockout miCLIP dataset. Here, we observed a complete loss of this peak, further confirming that this site is m^6^Am (Figure S3A).

Notably, the role of m^6^A in controlling stress-induced *Atf4* translation was described in mouse embryonic fibroblast cells (Zhou et al., 2018), rather than the HEK293T cells used here. Human cells appear to have lost the DRACH consensus sequence surrounding the putative m^6^A site (Figure S3B). Thus, it is possible that human cells exhibit stress-induced regulation of ATF4 translation through an m^6^A-independent pathway and mouse cells utilize an m^6^A-dependent pathway. To examine this possibility, we mapped 6mA in mouse embryonic fibroblasts using miCLIP (Figure S3C). Again, the 6mA peak was at the transcription-start site, not at a position corresponding to the second uORF (Figure S3D). These data further support the idea this peak derives from a m^6^Am residue. In comparison, there were low levels of 6mA reads throughout the transcript body suggesting either background reads or low stoichiometry m^6^A sites (Figure S3D). Thus, a role for a 5′ UTR m^6^A in uORF2 in regulating *ATF4* translation seems unlikely since the primary 6mA peak in *ATF4* is due to m^6^Am.

Overall, these data show that *PCIF1* deletion can be used to confirm whether a site is m^6^A or m^6^Am.

### Identification of internal 6mA sites that reflect m^6^Am rather than m^6^A

We noticed two unusual features in our mapping results. First, not all m^6^Am sites mapped to regions within annotated mRNA transcripts. Second, the m^6^Am metagene showed that while 94% of m^6^Am sites were located in the 5’ UTR, many were not directly at the annotated start sites and, in some cases, further downstream within the transcript body (Figure 3E).

We considered that these findings could be due to m^6^Am that occurs in mRNA isoforms that differ from the annotated transcripts due to an alternate transcription-start site. In the first case, m^6^Am could be at a transcription-start upstream of the transcription-start site in the RefSeq-annotated transcript, and therefore the m^6^Am is not assigned within a mRNA transcript. To test this, we plotted an m^6^Am metaplot relative to RefSeq-annotated transcription-start sites **(Figure 4A)**. Here we observed 16.7% of m^6^Am sites mapping within 250 nucleotides upstream of annotated start sites, suggesting that some m^6^Am occurs in isoforms with upstream start sites. We similarly observed m^6^Am upstream of transcription-start sites using GENCODE (Frankish et al., 2018) transcript annotations (Figure 4B). Using the FANTOM5 promoter-level expression atlas (Abugessaisa et al., 2017), a set of transcription-start sites specifically mapped across multiple tissues using the cap analysis gene expression (CAGE) approach, there was a marked overlap with our m^6^Am sites supporting the idea that these m^6^Am sites are indeed transcription-start sites (Figure 4C).

**Figure 4.**
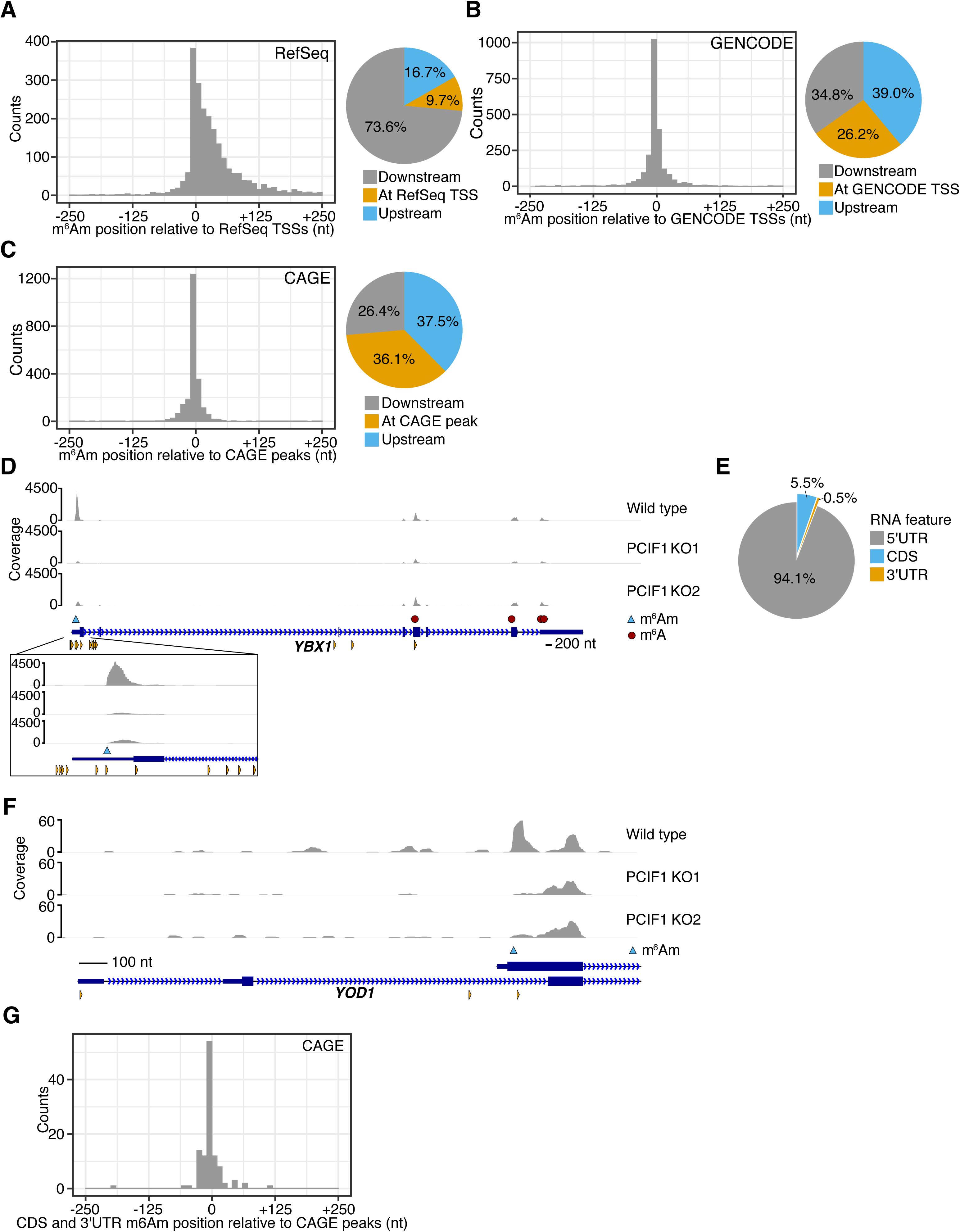
Internally mapped m^6^Am sites reflect m^6^Am in mRNA isoforms with alternative transcription-start sites. **A.** A metaplot centered on the closest RefSeq transcription-start site for each called m^6^Am site shows most m^6^Am sites are found downstream of the annotated start site, but not at the annotated start site. The distance of each m^6^Am to the nearest RefSeq transcription-start site (TSS, within 250 bp either side) was plotted as a histogram. The proportion of m^6^Am directly at the annotated transcription-start site, or up- or downstream, was plotted as a piechart, right. **B.** A metaplot analysis of m^6^Am locations using GENCODE transcription-start site annotations shows higher overlap with transcription-start sites. GENCODE annotations include more transcript isoforms and transcription-start sites than RefSeq. **C.** A metaplot of the distance from each m^6^Am site to the closest CAGE peak in the FANTOM5 database shows that m^6^Am sites are indeed transcription-start sites. To obtain a more comprehensive list of transcription-start sites, we used a CAGE dataset. Here, the overlap of m^6^Am was highest, suggesting that m^6^Am sites are selectively localized to transcription-start sites and not internal nucleotides within mRNA. **D.** The m^6^Am mapping to the annotated 5′ UTR of the YBX1 transcript reflects a transcript isoform. As shown, the PCIF1-dependent 6mA peak in YBX1 maps to the annotated 5′ UTR of *YBX1*. However, this peak overlaps with a CAGE site (orange triangles), indicating the existence of transcript isoform that initiates at the 6mA site. This further reinforces the idea that m^6^Am peaks that appear to be within the 5′ UTR instead reflect m^6^Am in transcript isoforms with alternate transcription-start sites. The exact m^6^Am nucleotide (blue triangle) was determined using the A to T transition within the PCIF1-dependent peak. Scale bar is shown. **E.** Most m^6^Am are found in the annotated 5′ UTR of transcripts. **F.** The internally mapping m^6^Am in *YOD1* derives from a transcript isoform. The m^6^Am peak in *YOD1* begins beyond the start codon of both annotated isoforms. Close-by CAGE peaks (orange triangles) suggest this is indeed a transcription-start site. **G.** A metaplot analysis of CDS and 3′ UTR mapping m^6^Am sites show the overlap with CAGE data and are thus transcription-start sites. The closest CAGE peak to each of the 6% of sites that appeared to not map to the 5′ UTR (E) was calculated and these distances were plotted as a histogram as in (C).

Transcription-start site heterogeneity likely explains why some m^6^Am sites map within the 5′ UTR, rather than solely located at the annotated transcription-start site. In the case of *YBX1*, a 6mA peak is mapped to the 5′ UTR and is lost in the *PCIF1* knockout miCLIP dataset, suggesting that this peak is due to m^6^Am (Figure 4D). This m^6^Am likely reflects an isoform with a transcription-start site located at this m^6^Am site, based on its overlap with a CAGE peak and PCIF1’s enzymatic preference for m^7^G capped mRNA (Figure 4D). Thus, the presence of m^6^Am within the 5′ UTR is likely to be a reflection of transcription-start site heterogeneity rather than “internal” m^6^Am nucleotides.

We next wanted to understand the basis for the ∼6% of m^6^Am sites that appear to map to coding sequences or 3′ UTR regions (Figure 4E). For example, *YOD1* shows an internal m^6^A peak in the first exon that is lost in the *PCIF1* knockout miCLIP dataset (Figure 4F). To determine if this m^6^Am site reflects a transcript isoform we examined the CAGE data for this transcript (Figure 4F). As with *YBX1* we similarly observed a transcription-start site that overlapped with the m^6^Am site. Thus, this internal site, which would normally have been assumed to be m^6^A using MeRIP-Seq and possibly miCLIP, derives from an isoform starting with m^6^Am.

To test this idea further, we performed a metagene analysis on m^6^Am sites mapping to coding sequences or the 3′ UTR. Here we plotted the distance to the nearest CAGE sites (Figure 4G). This analysis shows that many m^6^Am sites in the coding sequence and 3′ UTR are located at or near mapped CAGE sites. These data further suggest that m^6^Am is not internally located within transcripts but are instead found at the transcription-start sites.

Approximately 8% of m^6^Am sites that mapped to the coding sequence or 3′ UTR also contained an adjacent C to T transition. As a result, these peaks would likely have been called as an m^6^A. These data highlight the value of using *PCIF1* depletion to validate the transcriptome-wide m^6^Am and m^6^A maps.

### m^6^Am correlates with enhanced translation, expression, and stability of mRNAs

In our previous studies, we found that m^6^Am is correlated with transcripts that are highly expressed in cells (Mauer et al., 2017). We therefore wanted to reexamine this correlation based on the high-confidence m^6^Am annotation based on peaks that were depleted in the *PCIF1* knockout miCLIP dataset. In some cases, mRNAs that had been previously annotated as beginning with Am, Cm, Gm, or Um were re-annotated as m^6^Am for this analysis, and mRNAs previously annotated as m^6^Am were re-annotated as Am based on our revised mapping data. Analysis of mRNA expression showed that transcripts that begin with m^6^Am are indeed more highly expressed than mRNAs with other start nucleotides. Notably, among the highest expressed mRNAs in cells, m^6^Am appears to be the predominant start nucleotide (Figure 5A,B).

**Figure 5.**
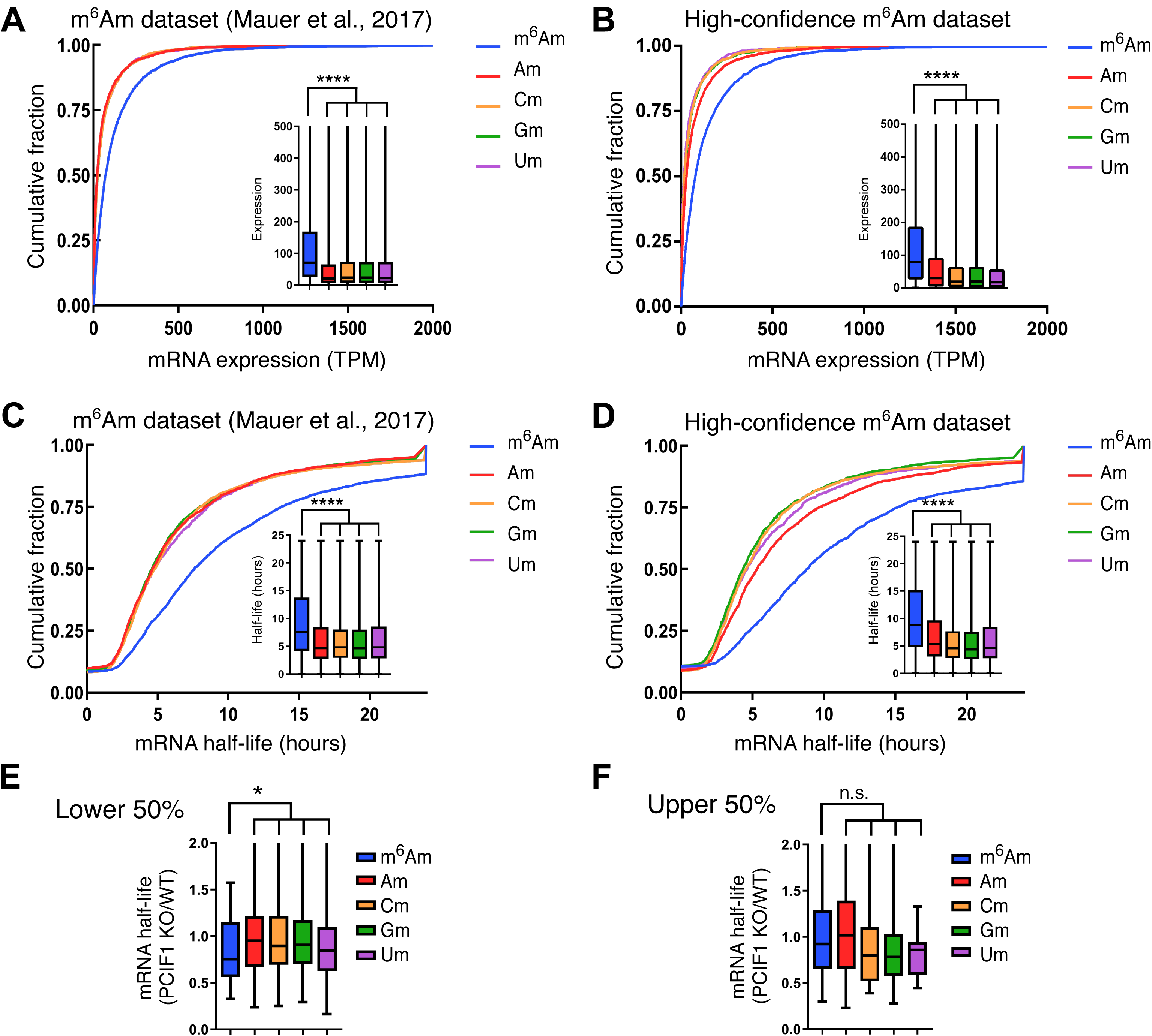
mRNA expression level and mRNA half-life depending on transcription start site. **A.** mRNAs with an annotated m^6^Am start nucleotide show higher mRNA expression than other mRNAs. mRNA expression level in wild-type HEK293T cells was based on the first annotated nucleotide and an earlier m^6^Am map (Mauer et al., 2017). Transcripts that start with m^6^Am are significantly upregulated. ****, P < 2.2 × 10–16, Student’s t-test. Cumulative distribution plot and boxplot represent the expression for mRNAs starting with m^6^Am, Am, Cm, Gm and Um. Data shown are the average gene expression measured from two replicates for HEK293T cells. **B.** m^6^Am mRNAs annotated using the *PCIF1* knockout miCLIP dataset show increased expression compared to mRNAs with other start nucleotides. Cumulative distribution plots were prepared as in **A** using the high-confidence m^6^Am dataset. The transcripts start with m^6^Am are significantly upregulated similarly as in **A**. ****, P < 2.2 × 10–16, Student’s t-test. Cumulative distribution plot and boxplot show the expression for mRNAs starting with m^6^Am, Am, Cm, Gm and Um. **C.** mRNAs with an annotated m^6^Am start nucleotide show higher mRNA half-life than other mRNAs. Annotated mRNA half-lives were based on the first annotated nucleotide and an earlier m^6^Am map (Mauer et al., 2017). mRNAs with an annotated m^6^Am exhibit a significantly elevated mRNA half-life than mRNAs with other annotated start nucleotides. ****, P ≤ 2.2 × 10–16, Student’s t-test. **D.** m^6^Am mRNAs annotated using the *PCIF1* knockout miCLIP dataset show increased expression compared to mRNAs with other start nucleotides. Cumulative distribution plot and boxplot of mRNA half-life are shown. Transcripts with m^6^Am have significantly longer half-life with similar P-value. ****, P ≤ 2.2 × 10-16, Student’s t-test. **E.** Influence of PCIF1 depletion of mRNA half-life for transcripts in the lower half of gene expression. Transcripts with m^6^Am have significantly shorter half-life in comparison to mRNAs annotated to begin with other nucleotides. *, P = 0.0258 by Student’s t-test. **F.** Influence of PCIF1 depletion on mRNA half-life for highly expressed transcripts. Transcripts with m^6^Am show no significant decrease in mRNA half-life in comparison to mRNAs annotated to begin with other nucleotides. n.s. (nonsignificant) by Student’s t-test.

We previously found that m^6^Am mRNAs show increased mRNA half-lives (Mauer et al., 2017) (Figure 5C). This effect was similarly seen with the high-confidence m^6^Am dataset (Figure 5D). Notably, for the mRNAs with an annotated half-life greater than 24 hours, the majority were m^6^Am transcripts.

Overall, these data suggest that the presence of m^6^Am correlates with an overall increase in mRNA stability, and that m^6^Am is the predominant starting nucleotide on “outlier” mRNAs with unusually high stability and expression.

We therefore wanted to understand the role of m^6^Am in these outlier mRNAs, as well as other mRNAs that use m^6^Am as the start nucleotide. We first we measured mRNA stability using SLAM-Seq (thiol(SH)-linked alkylation for the metabolic sequencing of RNA) in wild-type and *PCIF1* knockout HEK293T cells (Figure S4A). In this method, cells are pulsed with 4-thiouridine to enable incorporation in mRNA at approximately a 2% frequency relative to uridine (Herzog et al., 2017). Then, mRNAs are harvested at various points after chasing with uridine. The levels of 4-thiouridine are readily detected by treatment with iodoacetamide, which causes a U to C transition in RNA-Seq (Herzog et al., 2017).

To examine the outlier mRNAs, which are highly expressed, we separately examined mRNAs in the lower and upper half of gene expression. We only used transcripts that exhibited a minimum threshold of transitions required for mRNA half-life quantification. When we examined m^6^Am mRNAs in the lower half of gene expression, we observed a marked decrease in mRNA half-life upon PCIF1 depletion (Figure 5E). However, when we examined the more abundant mRNAs, which are enriched in the outlier transcripts, these transcripts did not show a substantial change in mRNA half-life (Figure 5F). We observed a slight reduction in stability relative to Am-annotated transcripts, but compared to all mRNAs (Am, Cm, Gm, and Um), these mRNAs appeared to show small, but nonsignificant increase in mRNA stability in *PCIF1* knockout cells. Thus, although m^6^Am is highly enriched in these outlier transcripts, the N6 methyl does not appear to account for their unusual stability. These transcripts may be unusually abundant as a result of a m^6^Am-independent mechanism. In contrast, mRNAs in the lower half of gene expression appear to utilize m^6^Am as part of their mechanism for transcript stability.

Previously we found that transcripts containing m^6^Am as the first nucleotide exhibit a subtle increase in translation relative to mRNAs with other start nucleotides (Mauer et al., 2017). To more directly test the role of m^6^Am on translation, we compared the translation efficiency of transcripts in control and *PCIF1* knockout cells by ribosome profiling (Figure S4B). Here, we found that transcripts that contained m^6^Am as the transcription-start nucleotide did not show a substantial change in translation efficiency upon PCIF1 depletion (Figure S4C). Rather than showing a decrease in translation, we observed a slight increase in translation upon loss of m^6^Am compared to transcripts annotated to begin with other nucleotides (Figure S4C).

Together, these experiments suggest that under the conditions used in these experiments, N6 methylation does not mediate the increase in translation efficiency of m^6^Am-initiated mRNAs in HEK293T cells.

## DISCUSSION

A major challenge when mapping m^6^A and m^6^Am is that both nucleotides are recognized by 6mA-specific antibodies and both can produce peaks in the 5′ UTR of mRNA transcripts. Although the miCLIP method helps to distinguish these modifications by detecting signature mutations and modification-specific peak shapes, these methods are not completely specific. As a result, determining whether an mRNA is regulated by m^6^A or m^6^Am can be challenging due to the inability to easily distinguish between these highly similar modifications. Here, by identifying PCIF1 and by subsequently depleting this m^6^Am-forming methyltransferase, we present a revised annotation of m^6^Am and m^6^A in the transcriptome. We find that previous annotation errors reflect the existence of mRNA isoforms that differ by transcription-start sites. In some cases, the isoforms contain transcription-start sites that map to internal sites within the annotated transcripts, resulting in the appearance of peaks that would otherwise be attributed to m^6^A. The identification and characterization of PCIF1 coupled with a precise m^6^Am annotations generated by PCIF1 depletion will facilitate the identification of functions for m^6^Am.

Using our new high-confidence m^6^Am map based on peaks that are lost in the *PCIF1* knockout miCLIP dataset, we find that m^6^Am is found on unusually stable transcripts that also exhibit increased transcript abundance in cells. Remarkably, depletion of m^6^Am by *PCIF1* knockout does not markedly impair the stability of these unusual transcripts under basal conditions. This suggests that these mRNAs utilize other mechanisms to achieve their unusual stability. m^6^Am may therefore have other functions in these transcripts which may only be revealed under specific cellular conditions.

However, m^6^Am has a clear stabilization effect on mRNAs that do not exhibit unusually high abundance. When mRNAs in the lower half of gene expression were examined, depletion of *PCIF1* lead to a marked reduction in the stability of m^6^Am-annotated transcripts. These mRNAs likely lack the specialized mechanisms that enable the unusually high expression of the outlier mRNAs. As a result, the stability of these mRNAs are sensitive to *PCIF1* depletion.

Other mechanisms that co-occur with m^6^Am formation likely account for the high stability of these unusual transcripts. Notably, the m^6^Am sequence motif resembles the YYANW motif, a 5-mer sequence corresponding to the transcription initiation site (initiator element; Inr) of RNA Polymerase II (Yang et al., 2007), suggesting that m^6^Am transcripts may preferentially derive from promoters containing this motif. It is intriguing to speculate that co-transcriptional mRNA processing events, other than m^6^Am formation, contribute to the unusual properties of these mRNAs.

In our previous studies, we found that m^6^Am was associated with transcripts that exhibit slightly higher translation levels than mRNAs that are annotated to begin with other nucleotides (Mauer et al., 2017). However, depletion of PCIF1 did not show an overall decrease in the translation efficiency of transcripts annotated to begin with m^6^Am. Thus, despite the clear difference if translation efficiency based on the presence of m^6^Am, it is likely that other mechanisms account for their increased translation.

A recent study by Akichika et al. used a previous m^6^Am annotation list and found that m^6^Am enhances translation based on analysis of *PCIF1* knockout cells (Akichika et al., 2018). However, the effect on mRNA translation upon *PCIF1* depletion was very small. Our analysis did not show an effect of m^6^Am on translation. Regardless, these two independent analyses suggest that the effects of m^6^Am on translation are likely to be small. It should be noted that effects of m^6^Am may be selectively seen during signaling or stress conditions that were not examined in either of these experiments.

Akichika et al. also found that *PCIF1* depletion did not affect the stability of m^6^Am mRNA (Akichika et al., 2018), which is consistent with our analysis of highly expressed m^6^Am-annotated mRNA. However, as we found, subsets of m^6^Am mRNAs may be preferentially regulated by m^6^Am in terms of mRNA stability, and potentially translation as well. Unlike m^6^A, which is largely found in one sequence context, m^6^Am can be followed by diverse nucleotides in mRNAs. It is likely that specific m^6^Am readers will be identified that are selective for m^6^Am based on its sequence context. Thus, the effects of m^6^Am are likely to be sequence and transcript specific.

A major goal of future studies will be to understand which cellular processes and pathways utilize m^6^Am. The identification of PCIF1 as the m^6^Am-forming methyltransferase will be an important step in understanding the physiological pathways regulated by this epitranscriptomic modification.

## ACKNOWLEDGMENTS

We thank J. Lipton for reagents and A. O. Olarerin-George for assistance with data analysis. This work was supported by NCN 2017/24/T/NZ1/00170 (D.T.S.), and NIH grants R00AG043550 and DP2AG055947 (E.L.G.), and R01DA037755 (S.R.J.).

## AUTHOR CONTRIBUTIONS

K.B. performed biochemical analysis of PCIF1 and generated PCIF1 knockout and overexpression cell lines, D.S. and K.B. performed assays of PCIF1 activity in cells, K.B., D.S., and S.Z. performed and analyzed ribosome profiling data, D.S. performed and analyzed SLAM-Seq experiments, B.H. performed and analyzed miCLIP experiments, N.L.I. performed cap-binding experiments, K.T. performed experiments assessing the translational effect of PCIF1 KO, T.G., J.-J. V and F.D. synthesized capped and uncapped RNA oligonucleotides, L.A.identified PCIF1 as putative m^6^Am methyltransferase, E.L.G and S.R.J. wrote the manuscript with input from all authors.

## METHODS

### Synthesis and characterization of synthetic oligonucleotides

The sequences of all the oligonucleotides used in this study are shown in Figure 1B.

The synthetic RNA oligonucleotides, used in Figure 1E, were chemically assembled on an ABI 394 DNA synthesizer (Applied Biosystems) from commercially available long chain alkylamine controlled-pore glass (LCAA-CPG) solid support with a pore size of 1000 Å derivatized through the succinyl linker with 5′-*O*-dimethoxytrityl-2′-*O*-Ac-uridine (Link Technologies). All RNA sequences were prepared using phosphoramidite chemistry at 1-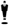 mol scale in Twist oligonucleotide synthesis columns (Glen Research) from commercially available 2′-*O*-pivaloyloxymethyl amidites (5′-*O*-DMTr-2′-*O*-PivOM-[U, C^Ac^, A^Pac^ or G^Pac^]-3′-*O*-(*O*-cyanoethyl-*N,N*-diisopropylphosphoramidite)(Lavergne et al., 2010) (Chemgenes). The 5′-terminal adenosine was methylated in 2′-OH (A_m_). The 5′-*O*-DMTr-2′-*O*-Me-A^Pac^-3′-*O*-(*O*-cyanoethyl-*N,N*-diisopropylphosphoramidite) (Chemgenes) was used to introduce A_m_ at the 5′-end of RNA. All oligoribonucleotides were synthesized using standard protocols for solid-phase RNA synthesis with the PivOM methodology(Lavergne et al., 2008).

After RNA assembly, the 5′-hydroxyl group of the 5′-terminal adenosine A_m_ of RNA sequences, still anchored to solid support, was phosphorylated and the resulting *H*-phosphonate derivative was oxidized and activated into a phosphoroimidazolidate derivative to react with either pyrophosphate (for ppp(A_m_)-RNA synthesis) (Zlatev et al., 2010) or guanosine diphosphate (for Gppp(A_m_)-RNA synthesis) (Thillier et al., 2012).

After deprotection and release from the solid support upon basic conditions (DBU then aqueous ammonia treatment for 4h at 37°C), all RNA sequences were purified by IEX-HPLC(Barral et al., 2013), they were obtained with high purity (>95 %) and they were unambiguously characterized by MALDI-TOF spectrometry.

*N*^7^-methylation of the purified Gppp(A_m_)-RNAs to give m^7^Gppp(A_m_)-RNAs was carried out quantitatively using human mRNA guanine-*N*^7^ methyltransferase and *S*-adenosylmethionine as previously described(Thillier et al., 2012). The oligonucleotides used in Figures 1C, 1D, 1G and 1H were synthesized by Trilink.

### Cell culture

HEK293T and HeLa cells were maintained in DMEM (11995-065, ThermoFisher Scientific) with 10% FBS and antibiotics (100 units/ml penicillin and 100 µg/ml of streptomycin) under standard tissue culture conditions. Cells were split using TrypLE^™^ Express (Life Technologies) according to the manufacturer’s instructions. Mycoplasma contamination in cells were routinely tested by Hoechst staining.

### Antibodies

Antibodies used for western blot analysis or immunostaining were as follows: mouse anti-FLAG M2 (F1804, Sigma, RRID: AB_262044), rabbit anti-PCIF1 (ab205016, Abcam, RRID: AB_2753142), mouse anti-β actin (A5441, Sigma, RRID: AB_476744), anti-eIF4E (2067, Cell Signaling, RRID: AB_2097675), anti-eIF4G (2498, Cell Signaling, RRID: AB_2096025). For m^6^A individual-nucleotide-resolution cross-linking and immunoprecipitation (miCLIP), rabbit anti-m^6^A (ab151230, Abcam, RRID: AB_2753144) was used.

### Generation of PCIF1 CRISPR knockout cells and overexpression cell lines

HEK293T and HeLa *PCIF1*-knockout cell lines were generated by CRISPR/Cas9 technology using two guide RNAs (gRNAs; 5′-CGGUUGAAAGACUCCCGUGG-3′ and 5′-ACUUAACAUAUCCUGCGGGG-3′) designed to target the PCIF1 genomic region between exon 8 and exon 17, that corresponds to the C terminal catalytic domain. Double-stranded DNA oligonucleotides corresponding to the gRNAs were inserted into the pSpCas9n(BB)-2A-Puro (PX459) V2.0 vector (62988, Addgene). Equal amounts of the two gRNA plasmids were mixed and transfected into HEK293T and HeLa cells using FuGENE 6 (Promega). The transfected cells were then subjected to puromycin selection for three days and viable cells were used for serial dilution to generate single-cell clones. The genomic deletion was screened by PCR and was confirmed by Sanger sequencing. HEK293T and HeLa *PCIF1*-knockout lines used in this study contained a 4655 or 4656 nt homozygous deletion that removed the region between exon 8 and exon 17, including the stop codon, resulting in the disruption of PCIF1 protein after P229 (aa 230-704). Loss of PCIF1 protein expression was confirmed by western blot with anti-PCIF1 antibody (Abcam).

Stable cell lines overexpressing PCIF1 WT or catalytically inactive mutant proteins were generated through retroviral infection. The coding sequence of human PCIF1 fused to a N-terminal 3X FLAG tag sequence that was cloned into the pBABE-puro retroviral vector (Addgene, 1764). Retroviral particles were generated in HEK293T cells through co-transfection of the packaging vectors pMD2.G (12259, Addgene) and pUMVC (8449, Addgene) with the appropriate pBABE-puro vectors. HEK293T and Hela cells were infected with retroviral particles of pBABE-puro-3X-FLAG-PCIF1 WT or pBABE-puro-3X-FLAG-PCIF1 SPPG or control pBABE-puro empty vector, followed by puromycin selection (1μg/ml).

Cells were maintained at 70-80% confluency before harvesting for mRNA purification. Two rounds of poly(A) mRNA isolation from mammalian cells was performed using oligo d(T)25 Magnetic mRNA isolation kit (NEB), according to the manufacturer’s instructions.

### Protein expression and purification

The coding sequence of human *PCIF1* was cloned as an in-frame fusion to the GST tagged vector pGEX-4T1. The catalytic site NPPF was mutated to APPA or SPPG thru site-directed mutagenesis using the Q5 mutagenesis kit (NEB), according to the manufacturer’s instructions. Recombinant GST-PCIF1 wild-type and catalytically inactive mutant proteins were expressed in *E. coli* T7 Express lysY. Overnight induction of protein expression was carried out with 0.5 mM IPTG at 18 °C. Bacteria were harvested at 4000 rpm, 4°C and the cell pellet was resuspended in protein purification lysis buffer (50 mM Tris-HCl pH 7.5, 0.25 M NaCl, 0.1% Triton-X, 1 mM PMSF, 1 mM DTT, and protease inhibitors). The lysate was sonicated 6 times in 30 seconds on/off cycles and then centrifuged at 12,000 rpm for 20 minutes. Lysates were incubated with glutathione Sepharose 4B beads (Sigma). Proteins and beads were washed 3 times with protein purification lysis buffer before incubating the beads with elution buffer (12 mg/ml Glutathione in protein purification lysis buffer, pH 8.0) for 30 minutes. Eluates were dialyzed overnight at 4 °C with enzyme storage buffer (40 mM Tris-HCl pH 8.0, 110 mM NaCl, 2.2 mM KCl, 1 mM DTT, 20% glycerol) and were subsequently stored at −80°C. Bradford assays and SDS-page gel electrophoresis followed by Coomassie staining was performed to determine integrity and quantity of purified proteins.

### *In Vitro* methyltransferase assays

*In vitro* methylation reactions (50 μl) assaying PCIF1 activity against the m^7^G capped RNA oligonucleotides were performed in methylation reaction buffer supplemented with 160 μM SAM (NEB) using 50 nM GST-PCIF1 protein and 4 μM m^7^G capped oligonucleotide. Reactions were incubated for 10 minutes at 37°C, followed by heat inactivation for 20 minutes at 65°C and subsequent clean up and buffer exchange using Biospin P6 columns (Biorad). RNA oligonucleotides were decapped using 25 Units of RppH (NEB) in ThermoPol buffer for 3 hours at 37°C, followed by clean up and buffer exchange with Biospin P6 columns. Decapped RNA oligonucleotides were digested to nucleosides with 2 units of Nuclease P1 (Wako USA) at 37°C for 3 hours in a buffer containing 10 mM ammonium acetate pH 5.3, 2mM ZnCl_2_ followed by treatment with 2 units of Fast Alkaline Phosphatase (FastAP, Thermo Scientific) in FastAP reaction buffer for 1 hour at 37°C. After digestion the sample volume was brought to 100 μl with ddH20 followed by filtration using 0.22 μm Millex Syringe Filters (EMD Millipore). 5 μl of the filtered solution was analyzed by UHPLC-MS/MS.

Enzyme kinetics assaying PCIF1 activity against the m^7^G-Am RNA oligonucleotide were performed in methylation reaction buffer supplemented with 1.33 μM [^3^H]-SAM (Perkin Elmer) and 10 μM SAM (NEB), using 20 nM GST-PCIF1 protein and a range of concentrations of m^7^G-Am oligonucleotide for 2-4 min at 37°C in 50 μl reactions. The reactions were stopped with 0.1% TFA followed by removal of unincorporated [^3^H]-SAM with Biospin P30 columns (Biorad). The purified RNA oligonucleotide samples were then subjected to scintillation counting using a Perkin Elmer scintillation counter. The Michaelis-Menten curve and K_M_ value were determined using Graphpad Prism software.

### UHPLC-ms/ms analysis

For the detection and quantification of internal m^6^A in mRNA, 500 ng of poly(A) mRNA was denatured at 70°C for 5 minutes followed by digestion to nucleotides using 20 units of S1 Nuclease (Thermo Scientific) in S1 Nuclease buffer for 2 hours at 37°C in 25 μl reactions. Nucleotides were then dephosphorylated to nucleosides by the addition of 2 units of Fast Alkaline Phosphatase (NEB) in FastAP reaction buffer for 1 hour at 37°C. After digestion the sample volume was brought to 100 μl with ddH20 followed by filtration using 0.22 μm Millex Syringe Filters (EMD Millipore). 5 μl of the filtered solution was analyzed by LC-MS/MS.

For the detection and quantification of cap-adjacent m^6^Am in mRNA, 500 ng of poly(A) mRNA was decapped using 25 Units of RppH (NEB) in ThermoPol buffer for 3 hours at 37 °C, followed by clean up and buffer exchange with Biospin P30 columns. Subsequently decapped RNA was denatured at 70 °C for 5 minutes followed by digestion to nucleotides using 2 units of Nuclease P1 (Wako USA) in a buffer containing 10 mM ammonium acetate pH 5.3, 2mM ZnCl_2_ for 3 hours at 37°C. Nucleotides were then dephosphorylated to nucleosides by the addition of 2 units of Fast Alkaline Phosphatase (NEB) in FastAP reaction buffer for 1 hour at 37°C. After digestion the sample volume was brought to 100 μl with ddH20 followed by filtration using 0.22 μm Millex Syringe Filters. 5 μl of the filtered solution was analyzed by LC-MS/MS.

The separation of nucleosides was performed using an Agilent 1290 UHPLC system with a C18 reversed-phase column (2.1 × 50 mm, 1.8 m). The mobile phase A was water with 0.1% (v/v) formic acid and mobile phase B was methanol with 0.1 % (v/v) formic acid. Online mass spectrometry detection was performed using an Agilent 6470 triple quadrupole mass spectrometer in positive electrospray ionization mode. Quantification of each nucleoside was accomplished in dynamic multiple reaction monitoring (dMRM) mode by monitoring the transitions of 268→136 (A), 282→136 (Am), 282→150 (m^6^A), 296→150 (m^6^Am), 244→112 (C). The amounts of A, C, Am, m^6^A and m^6^Am in the samples were quantified using corresponding calibration curves generated with pure standards. m^6^Am and m^6^A levels in the RNA oligonucleotides after *in vitro* methylation reactions were normalized by cytidine concentration. m^6^Am levels in mRNA were normalized by adenosine concentration.

### Cap-binding assay

Cells were lysed in buffer B (20 mM HEPES-KOH pH 7.6, 100 mM KCl, 0.5 mM EDTA, 0.4% NP-40, 20% glycerol) supplemented with protease and phosphatase inhibitors (Roche), 1 mM dithiothreitol (DTT) and 80 units/ml RNasin (Promega). For pull down, 1-2.5 mg of total protein extract was first pre-cleared on Agarose beads (Jena Bioscience) followed by incubation with 25 μl m^7^GTP conjugated Agarose beads (Jena Bioscience) for 1 hour at 4°C degrees. Following pull-down the beads were washed three times and the supernatant was removed and replaced by lysis buffer. Beads were incubated with 0.25 mM cap analog, m^7^GpppA, or GpppA, or water (mock) for 1 hour at 4 °C. Supernatant (Eluate) was removed and diluted with Laemmli sample buffer. Beads were washed three times and resuspended in Laemmli sample buffer. Samples were resolved on a 4–15% Tris-HCl gradient gel (BioRad) and analyzed by western blotting using specific antibodies.

### Immunofluorescence

Cells were grown on poly-L-lysine pre-coated coverslips that were sterilized under UV light for 30 minutes - 1 hour. Cells were rinsed in 1X phosphate-buffered saline (PBS) solution followed by fixation in ice-cold methanol at −20 °C for 10 minutes. Coverslips were then washed 3 times with 1X PBS before being blocked for 30 minutes in 1% BSA in 1X PBS. Primary antibody was diluted 1/200 in 1% BSA 1X PBS and incubated for 1 hour at room temperature in a humidified chamber. Slides were subsequently rinsed 3 times and washed 2 times for 15 minutes with 1% BSA in 1X PBS at room temperature before incubation with secondary antibody, diluted 1/200 in 1% BSA in 1X PBS, in a dark humidified chamber for 30 minutes at room temperature.

Coverslips were then rinsed 3 times and washed 3 times for 15 minutes with 1% BSA in 1X PBS in the dark before being rinsed 3 times with ddH_2_O. Coverslips were mounted using mounting medium containing DAPI. Image acquisition was carried out on a Nikon Eclipse Ti microscope (Nikon), using NIS-Elements AR software.

### Determination of relative m^6^A_m_, A_m_, and m^6^A levels by thin layer chromatography

Levels of internal m^6^A in mRNA were determined by 2D-TLC essentially as previously described (Zhong et al., 2008). In brief, poly(A) RNA (100 ng) was digested with 2 units ribonuclease T1 (ThermoFisher Scientific) for 2h at 37°C in the presence of RNasin RNase Inhibitor (Promega). T1 cuts after every guanosine and exposes the 5′-hydroxyl of the following nucleotide, which can be A, C, U, or m^6^A. Thus, this method quantifies m^6^A in a GA sequence context. 5′ ends were subsequently labeled with 10 units T4 PNK (NEB) and 0.4 mBq [γ-^32^P] ATP at 37°C for 30 min followed by removal of the γ-phosphate of ATP by incubation with 10 units Apyrase (NEB) at 30°C for 30 min. After phenol-chloroform extraction and ethanol precipitation, RNA samples were resuspended in 10 µl of DEPC-H_2_O and digested to single nucleotides with 2 units of P1 nuclease (Sigma) for 1h at 60°C. 1 µl of the released 5′ monophosphates from this digest were then analyzed by 2D-TLC on glass-backed PEI-cellulose plates (MerckMillipore) as described previously (Kruse et al., 2011).

The protocol to detect the m^6^A_m_:A_m_ ratio was based on the protocol developed by Fray and colleagues (Kruse et al., 2011), with some modifications. Poly(A) RNA (1 µg) was used for the assay. 300ng of poly(A) RNA was decapped with 15 units of RppH (NEB) for 3 h at 37°C. 5’ monophosphates in the resulting RNA were removed by addition of 5 units of rSAP phosphatase (NEB) for 1 h at 37°C. Up to this point, all enzymatic reactions were performed in the presence of SUPERase In RNase Inhibitor (ThermoFisher Scientific). After phenol-chloroform extraction and ethanol precipitation, RNA samples were resuspended in 10 µl of DEPC-H_2_O and 5′ ends were labeled using 30 units T4 PNK and 0.8 mBq [γ-^32^P] ATP at 37°C for 30 min. PNK was heat inactivated at 65°C for 20 min and the reaction was passed through a P-30 spin column (Bio-Rad) to remove unincorporated isotope. 8 µl of labeled RNA were then digested with 2 units of P1 nuclease (Sigma) for 1 h at 60°C. 2 µl of the released 5′ monophosphates from this digest were then analyzed by 2D-TLC on glass-backed PEI-cellulose plates (MerckMillipore) as described previously (Kruse et al., 2011).

Signal acquisition was carried out using a storage phosphor screen (GE Healthcare Life Sciences) at 200 µm resolution and ImageQuantTL software (GE Healthcare Life Sciences). Quantification was carried out with ImageJ (V2.0.0-rc-24/1.49m). For m^6^A_m_ experiments, the m^6^A_m_:A_m_ ratio was calculated. The use of this ratio has been described previously (Kruse et al., 2011). We confirmed that this assay is linear by spotting twice the sample material and confirming that the signal intensity doubles for the unmodified nucleotides (A, C, and U). Furthermore, exposure time of the TLC plates to the phosphor screen was chosen so the signal was not saturated. For m^6^A quantification, m^6^A was calculated as a percent of the total of the A, C, and U spots, as described previously (Jia et al., 2011). The use of relative ratios for each individual sample is important since it reduces the error derived from possible differences in loading. To minimize the effects of culturing conditions on the measured m^6^A_m_/A_m_ ratios of each experimental group (e.g. control vs. knockout), all replicates were processed in parallel to minimize any source of variability between samples being compared.

### miCLIP

Total RNA from wild-type and *PCIF1* knockout HEK293T cells, and wild type mouse embryonic fibroblasts, was extracted using TRIzol following the manufacturer’s protocol. Any contaminating genomic DNA was degraded using DNase I and poly(A) RNA was isolated using two rounds of Dynabeads Oligo(dT) capture. 10 µg poly(A) RNA was then used as input for single nucleotide-resolution m^6^A mapping using the miCLIP protocol, as previously reported (Linder et al., 2015). Final libraries were amplified and subjected to 50-cycle paired-end sequencing on an Illumina HiSeq2500 at the Weill Cornell Medicine Epigenetic Core facility.

### miCLIP bioinformatic analyses

The initial processing of raw FASTQ files was done as in the miCLIP protocol. Adapters and low quality nucleotides were first trimmed from paired reads using flexbar v2.5. The trimmed FASTQ file was then de-multiplexed using the pyBarcodeFilter.py script from the pyCRAC suite. The remainder of the random barcode was moved to the headers of the FASTQ reads using an awk script and PCR duplicates were removed using the pyCRAC pyDuplicateRemover.py script. Reads were aligned to hg38/mm10 using bwa v0.7.17 with the option “-n 0.06” as recommended in the CTK package. To identify m^6^A within the DRACH consensus, C to T transitions were extracted and the CIMS pipeline from the CTK package was used. Due to a high transition frequency in this dataset, putative m^6^A residues with an FDR<0.1 and in a DRACH consensus were used as the final list of m^6^A in this study.

To identify putative m^6^Am sites, coverage of wild type and *PCIF1* knockout samples were compared genome-wide using the bamCompare tool from deeptools v3.1.3. In short, the genome was binned into 50 nt non-sliding windows and the coverage of reads in each was counted for each strand, discarding zero-coverage bins. This was normalized to the total number of reads in bins, per million (BPM) and the log_2_ ratio of BPM+1 for wild type to *PCIF1* knockout was calculated. A log_2_ ratio threshold of 2 was chosen as the cutoff for each replicate. Adjacent bins passing threshold were merged using bedtools v2.27.1. The intersection of putative m^6^Am regions across replicates was taken using bedtools intersect, resulting in 2360 high-confidence m^6^Am peaks. To determine the precise m^6^Am nucleotide within these peaks, a combination of A to T transitions and a read pileup/drop-off method was used. In PCIF1-dependent peaks with an A to T transition occurring at a frequency of 10% or greater, this A was selected as the m^6^Am. For the remainder, a pileup/drop-off approach similar to the previous miCLIP criteria (Linder et al., 2015) was utilized. Here, the start nucleotide of each read (with respect to strand, i.e. the leftmost coordinate for + strand features and rightmost for – strand features) was extracted and piled up using the tag2cluster.pl script of the CTK package with the options “-s -v -maxgap −1”. Clusters of less than 5 reads were discarded, as were those that did not map to an A. When there was a single A-cluster in a PCIF1-dependent peak, this was selected as m^6^Am. When more than one occurred, the most piled-up cluster of the two closest to the beginning of the peak (with respect to strand) was selected.

To generate metagenes, MetaPlotR (Olarerin-George and Jaffrey, 2017). was used. In all case, the longest GENCODE transcript isoform for each gene was selected. For metaplots centered on reference annotations, the closest m^6^Am to each feature was measured using bedtools closest and these distances were plotted as a histogram. Aligned reads in bigwig format and BED files with coordinates for m^6^A, m^6^Am, and CAGE peaks were used to generate genome tracks using pyGenomeTracks v1.0. All motif searches were performed using DREME v5.0.2. For functional annotation analyses of m^6^Am and 5′ UTR m^6^A genes, DAVID v6.8 was used specifying a background of all genes covered with at least 20 reads.

### SLAM-seq

SLAM-seq was performed as described previously (Herzog et al., 2017) with minor modifications. HEK293T (WT and *PCIF1* KO) cells (at 60% confluency) were incubated with cell culture growth medium supplemented with 25 μM 4-thiouridine (s^4^U) for 24 h (pulse phase). s^4^U incorporation was confirmed by HPLC analysis, as previously described (Herzog et al., 2017). The uridine chase was initiated by changing media containing 2.5 mM uridine (Sigma) and cells were collected for RNA extraction after 6 and 12 h. The 0 h sample were the cells that have completed the pulse with s^4^U, but without uridine-chase. Total RNA was extracted using RNAzol reagent (MRC) according to the manufacturer’s instructions, maintaining reducing conditions to prevent oxidation of s^4^U (0.1 mM DTT final concentration). For thiol alkylation, a master mix (10 mM iodoacetamide, 50 mM NaPO_4_ pH 8 and 50% DMSO) was prepared, centrifuged, and added to 20 μg of total RNA at 50°C for 15 min and then purified by ethanol precipitation. After that, two rounds of poly(A) mRNA enrichment was carried out with oligo d(T)25 Magnetic Beads (NEB). Standard RNA-seq libraries were prepared using NEBNext Ultra Directional RNA Library Prep Kit for Illumina (NEB) following the instructions of the manufacturer. Sequencing was performed on a HiSeq2500 (Illumina) with 50 nucleotide reads.

### Ribosome profiling

Ribosome profiling was performed as described previously (McGlincy and Ingolia, 2017). In brief, wild-type and *PCIF1* knockout HEK293T cells were grown to ∼70% confluence, washed twice with ice cold PBS supplemented with 50 μg/ml of cycloheximide (CHX) and collected by scraping. After pelleting, cells were resuspended in 400 μl lysis buffer (20 mM Tris-HCl pH 7.4, 150 mM NaCl, 5 mM MgCl_2_, 1 mM DTT and 100 μg/ml CHX) After incubation on ice for 10 min, lysate was triturated 5 times through a 25-gauge needle and then lysate was centrifuged at 20,000 × g for 10 min. 5 μl of lysate was flash frozen and saved as input. To generate ribosome-protected fragments the lysates (30 µg) were first mixed with 200 μl DEPC-H_2_O then incubated with 15 U RNase I for 45 min at room temperature. The reaction was stopped with 10 μl SUPERase*In RNase inhibitor. 0.9 ml of sucrose-supplemented polysome buffer was added to the digestion mixture and ultracentrifuged at 100,000 rpm, 4°C for 1 h. Pellets were resuspended in 300 μl of water and after phenol-chloroform extraction, precipitated with ethanol. The RNA was then run on a 15% 8 M urea TBE gel, stained with SYBR Gold, and a gel fragment between 17-34 nucleotides corresponding to ribosome-protected RNA was excised. RNA was eluted for 2 h at 37°C in 300 μl RNA extraction buffer (300 mM NaOAc pH 5.5, 1 mM EDTA, 0.25%v/v SDS) after crushing the gel fragment. RNA was ethanol precipitated and resuspended in 26 μl water and treated with RiboZero Gold kit. Libraries from RNA-protected fragments were generated as previously described in the protocol (Linder et al., 2015). In brief, the RNA fragments were dephosphorylated with T4 PNK for 1 h at 37°C in dephosphorylation buffer (70 mM Tris, pH 6.5, 10 mM MgCl_2_, 1 mM DTT). The 3′ adaptor was ligated using T4 RNA Ligase 2, truncated K227Q ligase (New England BioLabs) for 3h at 22°C. Ligated sRNAs were purified by ethanol precipitation, and reverse transcribed using the primers complementary to the 3′ adaptor containing specific barcodes. After circularization with CircLigase II, cDNAs were relinearized by BamHI digestion and in the next step, PCR-amplified and subjected to Illumina HiSeq 2500 platform. Due to the similarity in size between ligated and unligated adapters, the libraries were gel purified.

RNA-Seq analysis was conducted using the ribosome profiling input material. Ribosomal RNAs were removed from the input RNA using the NEBNext rRNA Depletion Kit (NEB). Input RNA libraries were generated using the NEBNext Ultra Directional RNA library prep kit for Illumina (NEB). Libraries were sequenced using an Illumina HiSeq 2500 platform with 50 nt reads.

Ribosome footprint reads and corresponding RNA-Seq reads were processed essentially as described (Ingolia et al., 2012). Adaptors and short reads (<17nt) were trimmed using FLEXBAR v2.5, demultiplexed using pyBarcodeFilter.py (pyCRAC software). PCR duplicates were collapsed by pyFastqDuplicateRemover.py script. Ribosomal RNA reads were removed by STAR aligner38. Remaining reads were then aligned to the hg38 genome with STAR v2.5.2a in a splicing-aware manner and using UCSC refSeq as a transcript model database (version from June 02/2014 downloaded from Illumina iGenomes). Two mismatches were allowed and only unique alignments were reported. Aligned reads were then counted on transcript regions using custom R scripts considering only transcripts with annotated 5′ and 3′ UTRs. Gene count tables generated from STAR were normalized using DESeq2 (R-Bioconductor). Translation efficiency was calculated using Riborex (Li et al., 2017), with pre-filtering for transcripts that had at least ten counted reads.

### SLAM-seq bioinformatic analysis

Raw sequencing data were trimmed of adapter sequences and filtered of reads with uncalled bases and reads < 17 nucleotides in length using Flexbar. Duplicate reads were further removed using pyFastqDuplicateRemover.py script and remaining reads were aligned to the human genome (GrCh38) using the STAR aligner.

To identify T→C conversions, aligned reads were analyzed using Rsamtools Pileup (version 1.27.16). This program was used to determine the frequency of each of the four nucleotides present in mapped reads at every genomic position with read coverage. After summation of all nucleotide mapped to each transcript, we selected only those with at least 100 T→C conversions at time point 0 h. Additionally, to select for those transcripts with a longer half-life, transcripts were filtered for those with at least 50 T→C conversions at time point 6h. The mRNA half-life for each transcript was calculated based on the equation:

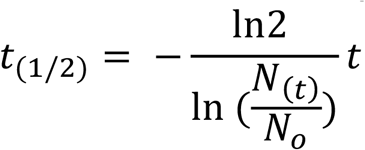

### Statistics and software

*P*-values were calculated with a two-tailed unpaired Student’s *t*-test or, for the comparison of more than two groups, with a one- or two-way ANOVA followed by Bonferroni’s or Tukey’s post-test. Reproducibility of half-life and translation efficiency measurements was assessed by calculating the Spearman correlation coefficient between replicates. Significance of list overlaps was calculated using hypergeometric probability.

## CONTACT FOR REAGENT AND RESOURCE SHARING

Please contact E.L.G. or S.R.J. for reagents and resources generated in this study.

## SUPPLEMENTARY FIGURE LEGENDS

**Figure S1.**
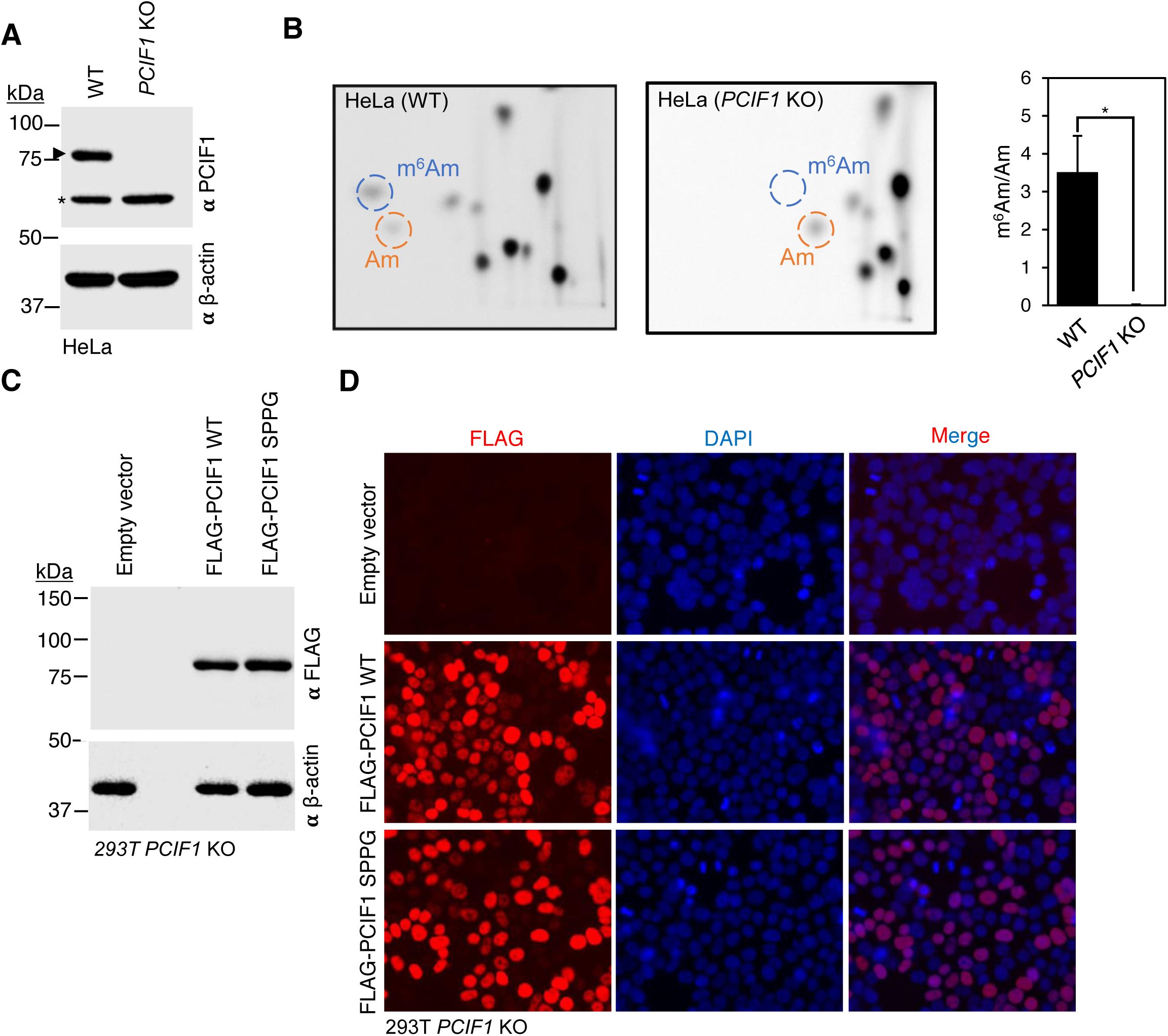
PCIF1 deletion eliminates m^6^Am in HeLa cells and both wildtype and catalytically dead PCIF1 express and localizes similarly. **A.** CRISPR-mediated knockout of *PCIF1* in HeLa cells is efficient as assessed by western blots of whole cell extracts blotted with an anti-PCIF1 antibody. The upper band represents endogenous PCIF1, whereas the lower band is a non-specific band. β-actin is shown as a loading control. **B.** m^6^Am is depleted in *PCIF1* KO HeLa cells as assessed by 2D-TLC. Left panels show representative images of 3 independent experiments with the blue circles representing m^6^Am and the orange circles representing Am as assessed by standards run on separate blots in parallel. The bar graph on the right represents the mean ± s.e.m of 3 independent experiments. *: p < 0.05 as assessed by ratio paired t-test. **C.** Western blot analysis demonstrate equivalent levels of wild-type and catalytically inactive PCIF1 rescue in HEK 293T *PCIF1* KO cells. Whole cell extracts from HEK293T *PCIF1* KO control cells (Empty Vector) and HEK 293T *PCIF1* KO cells stably overexpressing 3X-FLAG-PCIF1 WT or 3X-FLAG-PCIF1 SPPG catalytic mutant were blotted with anti-FLAG antibody (upper panel) and β-actin (lower panel). **D.** Immunofluorescence staining of PCIF1 rescue demonstrates equivalent levels and localization of wildtype and catalytically dead PCIF1 in HEK293T *PCIF1* KO cells stably expressing either 3X-FLAG-PCIF1 WT or 3X-FLAG-PCIF1 SPPG catalytic mutant. Both PCIF1 WT and SPPG mutant are expressed at similar levels and are primarily localized to the nucleus. DAPI was used to stain nuclei. Representative images are shown.

**Figure S2.**
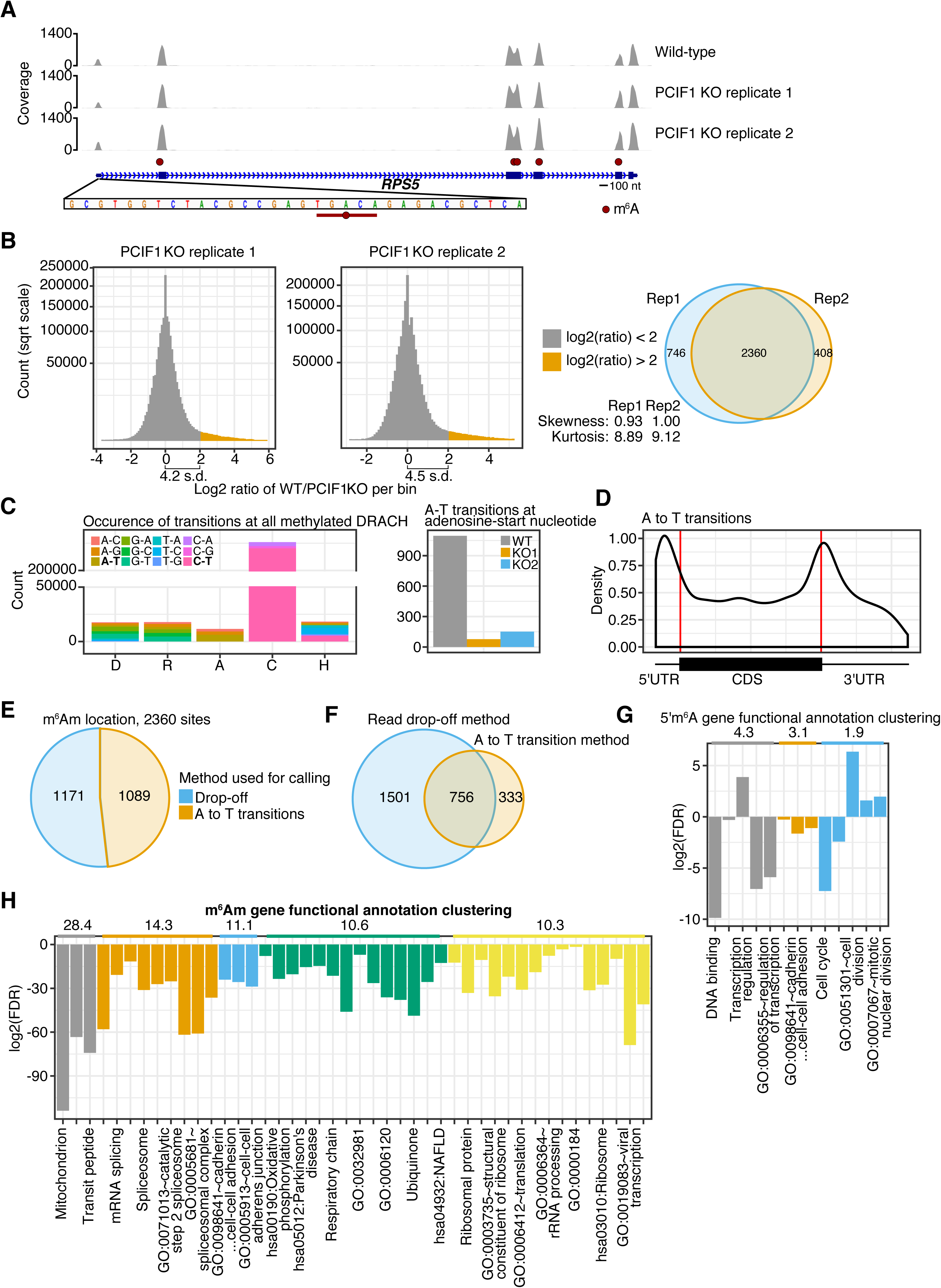
*PCIF1* KO and miCLIP A to T transitions can be used to identify high-confidence m^6^Am sites in the transcriptome. **A.** As in Figure 3D, *RPS5* is likely to be a 5′ UTR m^6^A-containing transcript instead of m^6^Am, as was previously thought. The persistence of the transcription-start site-proximal peak in the *PCIF1* knockouts suggests that this must be dependent on m^6^A. No m^6^A (as called by C to T transitions) was detectable at this peak, which may be due to the low and stochastic nature of C to T transitions. Inset shows the nucleotide sequence at this peak at the start site, with a potential DRACH motif (TGACA) indicated by the red line. Called m^6^A sites are shown as red circles, and scale bar is shown underneath the gene structure. **B.** Analysis of the fraction of miCLIP reads that are dependent on PCIF1. In this experiment, the genome was binned into 50 nt-wide windows and miCLIP read coverage within each bin was calculated with respect to strand. These values were normalized to the total number of reads mapping across all non-zero-coverage bins, per million (BPM). The log_2_ ratio of wild-type BPM+1 over knockout BPM+1 per was then plotted as a histogram with a square root y-axis. For each replicate, the skewness was ∼1 with a kurtosis of ∼9, showing that the very few deviations from no change were mainly found with a higher ratio. A cut off of 2 was chosen as being putative m^6^Am regions per replicate. This was over 4 standard deviations from the mean. Overlap analysis showed a highly similar set of m^6^Am regions. The intersection between the two replicates was then used as a list of high-confidence m^6^Am regions as shown in the Venn diagram. These regions were then used for m^6^Am site calling. **C.** A to T transitions are the most common transition at methylated adenosines and are markedly reduced in the *PCIF1* knockout miCLIP dataset. The transitions for each nucleotide at m^6^A-called DRACH sites was plotted (left). As expected, the major m^6^A signature mutation is the C to T transition as reported previously. However, at the m^6^A site, A to T transitions, although less common than C to T transitions, are the most common mutation. These A to T transitions are markedly reduced at m^6^Am sites in the *PCIF1* knockout (right). Thus, *PCIF1*-dependent A to T transitions can be used to confirm a m^6^Am site. **D.** Metagene analysis for all A to T transitions in miCLIP. A to T transitions are found primarily at the stop codon and TSS, likely reflecting antibody-induced signature mutations induced by m^6^A and m^6^Am, respectively. **E.** Piechart showing the methods used for m^6^Am site calling. Of the 2360 high-confidence m^6^Am regions, 1089 A to T transitions occurred at 10% or more reads. For those regions that did not meet these criteria, the most likely A-starting read pileup was chosen. **F.** Overlap in m^6^Am sites called by A to T transitions compared to the drop-off method. The A to T transitions provide high confidence that the chosen nucleotide is a 6mA; nevertheless, most sites were accurately called using the read-start method. **G.** Functional annotation clustering shows 5’UTR m^6^A is associated with transcription and the cell cycle. The non-redundant list of 5’UTR m^6^A was annotated using DAVID v2.8. A dataset-specific background of all genes with at least 20 reads was used for this analysis. DAVID clusters enriched and similar Gene Ontologies and other pathways so that an overview of the general cellular processes of the gene sets investigated are involved in can be determined. Each cluster is similarly colored; the enrichment score is shown above for each cluster. Every other term title is shown due to space restraints; these are available in **Table S3**. **H.** Function annotation clusters of m^6^Am genes show these are strongly associated with mRNA splicing, mitochondrial processes, and translation. DAVID function annotation was performed as in (G) using the list of high-confidence m^6^Am genes and the dataset-specific background. Clusters are shown in the same color and each clusters enrichment score is shown above. Every other term title is shown due to space restraints; these are available in **Table S3**.

**Figure S3.**
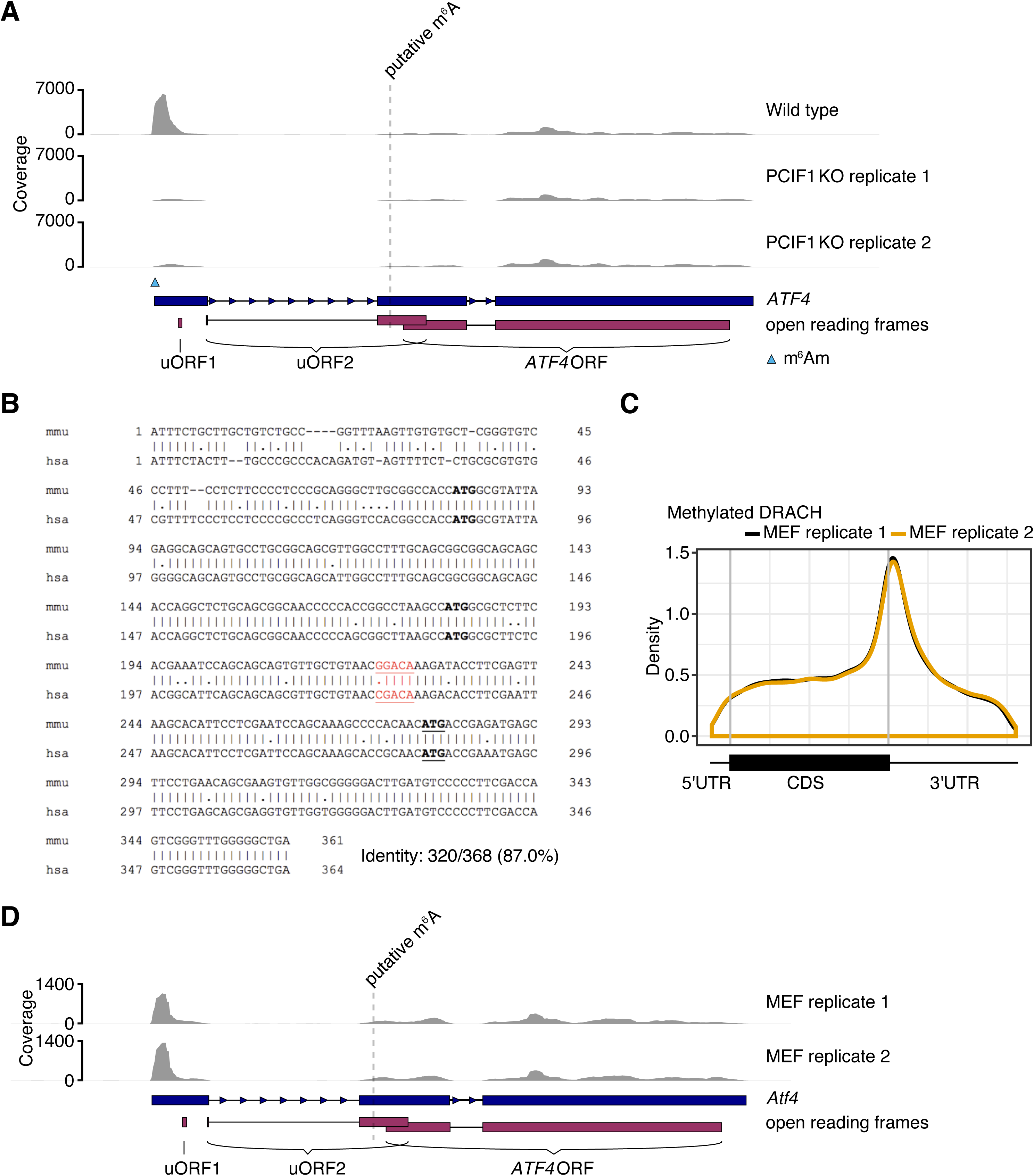
*ATF4* shows a PCIF1-regulated m^6^Am peak but no specific enrichment of m^6^A in uORF2 or elsewhere in the mRNA. **A.** The largest 6mA peak in *ATF4* is due to m^6^Am. The genome track for *ATF4* in wild-type HEK293T cells shows that the major peak is at the transcription-start site. This site is lost in the *PCIF1* knockout cells. Shown is the location of the putative m^6^A site in uORF2 as was reported in (Zhou et al., 2018). The schematic below indicates the complex uORF structure of the *ATF4* mRNA. **B.** Pairwise alignment of the human and mouse *ATF4* sequences show that the putative m^6^A DRACH consensus site is only seen in mouse *ATF4*, but not human *ATF4*. The mouse and human sequences from the transcription-start site to the end of uORF2 (which extends beyond the start codon of the coding sequence of the main *ATF4* ORF) were aligned. The start codons for uORF1, uORF2, and *ATF4* coding sequence are indicated in bold and underlined. The uORF sequences share high (87%) sequence identity; however, at the putative mouse m^6^A site, the DRACH consensus was lost (GGACA in mouse to CGACA in humans, shown in red). **C.** miCLIP analysis of mouse embryonic fibroblast poly(A) RNA. The miCLIP analysis shows the expected stop codon enrichment of m^6^A. The called m^6^A sites were plotted as a metagene using MetaPlotR. Both MEF replicates show the typical m^6^A metagene with a high degree of overlap. **D.** A genome track of mouse *Atf4* (NM_009716.1; (Vattem and Wek, 2004)) using the MEF miCLIP data show a very similar profile to the human miCLIP profile on *ATF4* shown in (A). The major 5′ UTR 6mA peak is similarly enriched at the transcription-start site, with little to no reads at the putative m^6^A site. Due to the highly similar profile, and the high identity of the mouse and human 5′ UTR sequence (Supplementary Figure 4B), the major 6mA peak in MEFs is most likely m^6^Am. m^6^A is unlikely to be present within the body of this transcript except at trace levels.

**Figure S4.**
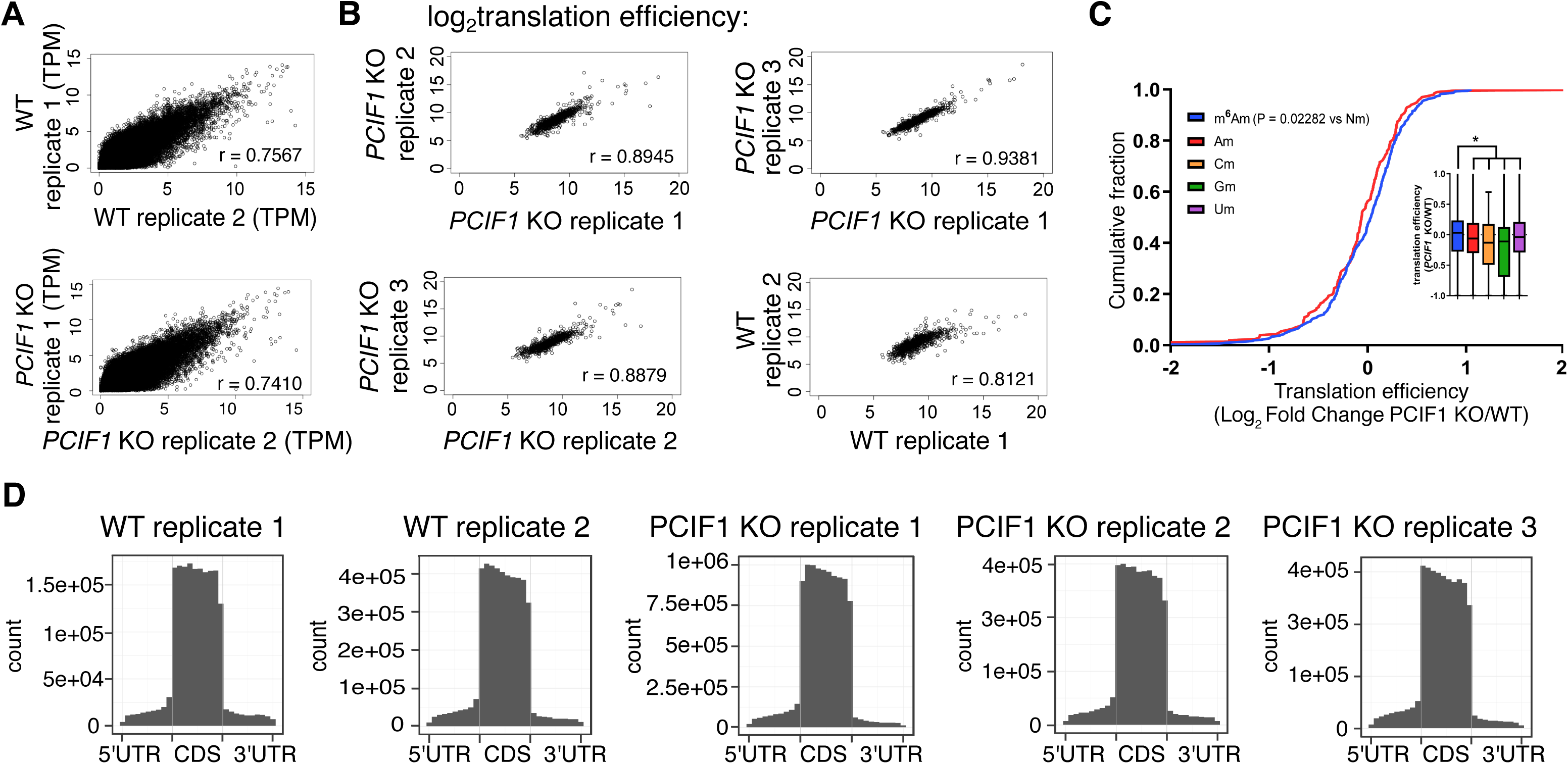
m^6^Am deficiency influences translation efficiency. **A.** Correlation of Slam-Seq replicates derived from HEK293T wild-type and *PCIF1* knockout cells (0 h). The Spearman correlation coefficient (r) is shown **B.** Ribosome profiling dataset replicates show high correlation. Shown are the correlation plots of translation efficiency for two wild-type HEK293T replicates and three *PCIF1* knockout HEK293T replicates. The Spearman correlation coefficient (r) is shown. **C.** Effect of PCIF1 depletion of mRNA translation efficiency. Ribosome profiling was performed in wild-type and *PCIF1* knockout HEK293T cells. A matched RNA-Seq experiment was performed for each replicate in order to calculate translation efficiency. The boxplot represents the change in translation efficiency for mRNAs based on their annotated start nucleotide. The cumulative distribution plot indicates the change in translation efficiency in the *PCIF1* KO relative to wild-type HEK293T cells for mRNAs with an annotated m^6^Am start nucleotide in comparison to mRNAs with an annotated Am start nucleotide. The translation efficiency of mRNAs starting with an m^6^Am is slightly increased upon *PCIF1* KO. Data represent the average from two independent replicates of ribosome profiling data sets for wild-type HEK293T cells and three replicates of *PCIF1* KO HEK293T cells. *, *P* = 0.02282 by Student’s t-test. **D.** Ribosome-protected fragments are enriched in the coding sequence for all replicates. Shown is the distribution of ribosome profiling reads between the coding sequence (CDS) and UTRs. These data show the expected high coverage in the CDS compared to UTRs.

